# A fully automated FAIMS-DIA proteomic pipeline for high-throughput characterization of iPSC-derived neurons

**DOI:** 10.1101/2021.11.24.469921

**Authors:** Luke Reilly, Lirong Peng, Erika Lara, Daniel Ramos, Michael Fernandopulle, Caroline B. Pantazis, Julia Stadler, Marianita Santiana, Anant Dadu, James Iben, Faraz Faghri, Mike A. Nalls, Steven L. Coon, Priyanka Narayan, Andrew B. Singleton, Mark R. Cookson, Michael E. Ward, Yue A. Qi

## Abstract

Fully automated proteomic pipelines have the potential to achieve deep coverage of cellular proteomes with high throughput and scalability. However, it is important to evaluate performance, including both reproducibility and ability to provide meaningful levels of biological insight. Here, we present an approach combining high field asymmetric waveform ion mobility spectrometer (FAIMS) interface and data independent acquisition (DIA) proteomics approach developed as part of the induced pluripotent stem cell (iPSC) Neurodegenerative Disease Initiative (iNDI), a large-scale effort to understand how inherited diseases may manifest in neuronal cells. Our FAIMS-DIA approach identified more than 8000 proteins per mass spectrometry (MS) acquisition as well as superior total identification, reproducibility, and accuracy compared to other existing DIA methods. Next, we applied this approach to perform a longitudinal proteomic profiling of the differentiation of iPSC-derived neurons from the KOLF2.1J parental line used in iNDI. This analysis demonstrated a steady increase in expression of mature cortical neuron markers over the course of neuron differentiation. We validated the performance of our proteomics pipeline by comparing it to single cell RNA-Seq datasets obtained in parallel, confirming expression of key markers and cell type annotations. An interactive webapp of this temporal data is available for aligned-UMAP visualization and data browsing (https://share.streamlit.io/anant-droid/singlecellumap). In summary, we report an extensively optimized and validated proteomic pipeline that will be suitable for large-scale studies such as iNDI.

## Introduction

The human induced pluripotent stem cell (iPSC) Neurodegenerative Disease Initiative (iNDI), a program within the NIH Center for Alzheimer’s and Related Dementias (CARD), was established to apply genome-engineered iPSC-derived neuron models to the field of Alzheimer’s disease and related dementias (ADRD) research (Ramos et al., 2021). iNDI aims to comprehensively characterize isogenic iPSC lines harboring 134 variants across 73 ADRD genes using proteomics, transcriptomics, high-throughput microscopy, and CRISPRi-based screens. Over time, iNDI will generate a large panel of ADRD-relevant iPSCs and associated phenotypic data that will be openly shared with the larger scientific community. Long-term, large-scale projects require well-validated pipelines to counter risks of outcome reliability and consistency. Here, we report an optimized data acquisition and database search strategy for a mass spectrometry (MS)- based whole-cell, single-shot proteomics approach for iNDI, which prioritizes proteome coverage, reproducibility, accuracy, and throughput.

A major challenge in the study of neurodegenerative disorders is creating suitable *in vitro* models of disease with human cells. Since mature neurons are post-mitotically inert, they are difficult to manipulate genetically (Frade & Ovejero-Benito, 2015). In contrast, iPSC-derived neurons represent a tractable model for genomic editing to study ADRD genes and their variants. A thorough characterization of iPSCs and differentiated neurons is fundamental to developing baseline data for subsequent analyses of iPSC-derived models of disease.

Traditional mass spectrometric analysis typically employs a data dependent acquisition (DDA) mode, which selectively performs fragmentation scans (i.e., MS2) on the highest intensity precursor ions (i.e., MS1). This approach typically results in missing values and poor reproducibility, which is especially problematic in large-scale studies. In contrast, data independent acquisition (DIA), in which all precursors are fragmented regardless of intensity, significantly improves both throughput and reproducibility (Muntel et al., 2019). Due to its MS2 transition-based quantification, DIA can provide higher accuracy than the MS1-based DDA method (Shi et al., 2016). With rapidly advancing scan speed, sensitivity, and resolution in next generation mass spectrometer systems, single-shot analysis has become a common approach to maximize reproducibility while minimizing sample processing times. DDA-based single-shot proteomics often results in lower proteome depth compared to fractionated approaches (Shishkova et al., 2016); in contrast, DIA-based single-shot proteomics achieves greater proteome depth (Muntel et al., 2019).

Prior to mass-spectrometry, complex samples typically require separation that is achieved through the use of a variety of approaches, such as offline liquid phase fractionation, SDS-PAGE and ion mobilty interface. Of these, high-field asymmetric ion mobility spectrometer (FAIMS) interface separates ion groups within an electric field with applied compensation voltages (CVs), thereby determining which ions pass through the device. Within the DDA paradigm, FAIMS has been shown to enhance protein identification at the expense of reduced peptide coverage (Bekker-Jensen, Martinez-Val, et al., 2020; Hebert et al., 2018). However, single-shot FAIMS-assisted DIA has been relatively unexplored. Therefore, we systematically evaluated and optimized FAIMS-DIA in conjunction with applied database search strategies.

Here, we performed proteomic profiling of KOLF2.1J over the course of a 28-day neuron differentiation using the optimized FAIMS-based single-shot DIA approach. This study demonstrates the strength of FAIMS-DIA for proteome analysis and provides robust datasets detailing changes in protein abundance throughout differentiation from iPSC to neurons. These data provide a framework to assess neuronal protein expression during differentiation and maturation and the relative abundance of ADRD genes in iPSCs compared to differentiated neurons.

## Methods

### Human iPSCs differentiation

We adopted the neuron differentiation protocol described previously (Fernandopulle et al., 2018). This protocol involves overexpression of a doxycycline-inducible human neurogenin-2 (NGN2), with the modification in this study that NGN2 was delivered via a PiggyBac Transposase system. KOLF2.1J and WTC11 (i.e. i11w-mNC)(Tian et al., 2019) iPSCs were transfected with both a PB-TO-hNGN2 vector (Addgene plasmid #172115) and EFa1-Transposase (Addgene plasmid #172116) in a 1:2 ratio (transposase:vector) using Lipofectamine (Thermo Scientific Cat. #STEM00015), followed by selection for 2 weeks with 8 μg/mL of puromycin (Sigma-Aldrich Cat. #P9620).

Post-selection iPSCs with stably-integrated human NGN2 under a tetracycline-inducible promoter were dissociated into single cells using Accutase (Thermo Scientific Cat. #A1110501) and plated on a Matrigel (1:100, Corning Cat. #354277)-coated 6 well, with 1.5 million cells per well. After this plating step (day 0 of protocol), cells were cultured for 3 days in induction media (IM) consisting of Knockout DMEM/F12 (Thermo Scientific Cat. #12660012), 1X N-2 Supplement (Thermo Scientific Cat. #17502001), 1X Non-Essential Amino Acids (Thermo Scientific Cat. #11140050), and 1X Glutamax (Thermo Scientific Cat. #35050061), with 10 μM of ROCK inhibitor Y-27632 (SelleckChem Cat. #S1049) and 2 μg/mL Doxycycline (Sigma-Aldrich Cat. #D9891) added the day of. On day 3, cells were again dissociated with Accutase and re-plated into a 96-well plate coated with poly-L-ornithine hydrobromide (PLO, 0.1 mg/mL, Sigma-Aldrich Cat. #P3655), with 30,000 cells per well. The induced cells were then differentiated for 25 days using Brainphys (Stemcell Technologies Cat. #5790) neuronal maturation medium containing 1X B-27™ Plus Supplement (Thermo Scientific Cat. #A3582801), 10 ng/mL GDNF (PeproTech Cat. #450-10), 10 ng/mL BDNF (PeproTech Cat. #450-02), 10ng/mL NT3 (PeproTech Cat. #450-03), 1μg/mL Laminin (Trevigen Cat #3446-005-01) and 2 μg/mL Doxycycline (Sigma-Aldrich Cat. #D9891). For neuronal maintenance, half the media was changed every 3-4 days. For neurite outgrowth, we stably expressed cytosolic mScarlet (MK-EF1a-mScarlet)(Tian et al., 2019). The iPSCs with a stably-integrated human NGN2 were then transduced with lentivirus expressing cytosolic mScarlet to identify neurites.

### Imaging and Image analysis

The course of the neuron differentiation was recorded by scanning using an Incucyte® S3 Live-Cell Analysis System with a 20X objective. Imaging was performed every 24 h at 37°C for 25 days. Phase and fluorescent images were acquired for every time point. The neurite outgrowth was analyzed with NeuroTrack Incucyte software module.

### Fully automated sample preparation for proteomics

For the temporal proteomic profiling in KOLF2.1J-derived iNeurons, we used a fully automated sample preparation workflow (**Supplemental Fig. S1A**) based on the singlepot, solid-phase-enhanced sample preparation (SP3) approach (Hughes et al., 2019). Cells were washed thoroughly washed with ice-cold PBS (Lonza Cat. #17-516F/12) and 100 μL SP3 lysis buffer were directly added in each well of a 96-well plate without cell lifting (SP3 lysis buffer: 50 mM Tris-HCI [pH = 8.0], 50 mM NaCl, 1% SDS, 1% triton X-100 (MilliporeSigma Cat. #X100), 1% NP-40 (Thermo Scientific Cat. # 85124), 1% tween 20 (MilliporeSigma Cat. #P9416), 1% glycerol (MP Biomedicals Cat. #800687), 1% sodium deoxycholate (wt/vol) (MilliporeSigma Cat. # D6750), 5 mM EDTA [pH = 8.0], 5mM dithiothreitol [DTT] (Thermo Scientific Cat. # 20290), 5KU benzonase (MilliporeSigma Cat. #E8263), and 1× complete protease inhibitor (MilliporeSigma Cat. # 5892970001). Cells in the orginal 96-well tissue culture plate were lysed and reduced on a ThermoMixer pre-heated to 65 °C at 1200 r.p.m. for 30 minutes. Samples were subsequently alkylated through addition of iodoacetamide (Thermo Scientific Cat. #A39271) to a final concentration of 10 mM and shielded from light for 30 minutes.We performed an automated detergent compatible protein assay (DCA) (Bio-Rad, Hercules, CA, Cat. #5000111), normalization (to 1 μg/μL) and radomlization, and aliquoted 20 μg protein per well to KingFisher deep 96-well plates on an Agilent Bravo liquid handling platform. Then, we conducted an automated SP3 enrichment platform on a KingFisher APEX robot (Thermo Scientific). Briefly, cell lysate was pre-mixed with 100% ethanol at 1:1 volume ratio and was incubated with pre-washed SP3 beads (Millipore Sigma Cat. #45152105050250 and #65152105050250, 1:1 ratio, 50 μg/μL) for 10 minutes at a slow mixing rate; the beads with enriched proteins were washed three times with 80% ethanol (Electron Microscopy Sciences Cat. #15055) followed by incubation with 100 μL 50mM ammonium bicarbonate (MilliporeSigma Cat. #A6141) addition of 1 μg trypsin/lysC mix (Promega, Cat. # V5071) at 37 °C for 16 hours. The resulting tryptic peptides were freeze-dried and reconstituted in 2% acetonitrile (ACN) in 0.1% trifluoroacetic acid (TFA). Peptide concentration was evaluated via a Denovix® DS-11 FX (peptide mode, acquisition wavelength of 215 nm, E 0.1% (mg/mL) correction factor set to 25.99). Peptides were normalized to 0.2 μg/μL, and 5 μL of sample was used for each run, resulting in a total of 1 μg peptide analyzed by LC-MS/MS. The Hela Protein Digest Standards (ThermoFisher Cat. #88328) were directly reconstituted to 0.2 μg/μL using 2% ACN in 0.1% TFA.

### Liquid chromatography and mass spectrometry

An UltiMate™ 3000 nano-HPLC system coupled to a hybrid Orbitrap Eclipse™ mass spectrometer LC/MS system was used for all experiments. The nano-HPLC had identical settings for all LC-MS/MS runs to exclude the possibility of LC interference. Liquid phase A (5% DMSO in 0.1% formic acid (FA), in water) and liquid phase B (5% DMSO in 0.1% FA, in ACN) were used. The tryptic peptides were separated on a ES903 nano column (75 μm × 500 mm, 2 μm C_18_ particle) using a 2-hour efficient linear gradient with constant flow rate of 300 nL/min, 0 – 5 min (2% phase B, sample loading), 5 - 120 min (2% - 35% phase B), 120-125 min (35% −80% phase B), 125 – 135min (80% phase B), 135 – 136 min (80% - 2% phase B), 136 – 150 min (2% phase B). Samples were loaded on a trap column (75 μm × 20 mm, 3 μm C_18_ particle) using the loading pump at a constant rate of 5 μL/min. The nano and trap columns were heated to 60 °C in the EASY-Spray electronic ionization source and column oven, respectively.

DDA MS1 resolution, regardless of FAIMS usage, was set to 120K, scan range was 375-1400 m/z, and RF lens was 30%. The ions with charge state 2-4 were selected for MS2 fragmentation, dynamic exclusion was set to 45 s, and minimum intensity threshold was set to 50000. The cycle time of the data dependent mode was set to 3 s. For the MS2 acquisition, we used an isolation window of 1.6 m/z in quadrupole and ion trap coupled with HCD-enabled fragmentation with collision energy at 30%. The maximum injection time was set to 35 s, and AGC target was 100%.

For DIA experiments, regardless of FAIMS usage, MS1 resolution was set to 120K, and both standard AGC target and auto maximum injection time were selected. For MS2 scans, the precursor range was set to 400-1000 m/z, and the isolation window was 8 m/z with 1 m/z overlap, resulting in 75 windows for each scan cycle. The MS2 fragmentation was performed in Orbitrap with 30K resolution and HCD with 30% collision energy. The AGC target was set to 800%. The MS2 scan range was defined as 145-1450 m/z, and the loop control was set to 3 s **(Supplemental Fig. S1B)**.

### Liquid Phase Fractionation

For the offline liquid phase fractionation (LPF), HeLa digest was fractionated into 8 fractions using a high pH reversed-phase peptide fractionation Kit (ThermoFisher, Cat. #84868). Twenty μg of HeLa digest was loaded to a C_18_ solid phase spin column and eluted with a serial concentration of ACN (5% to 50%). Subsequently, the peptides from 8 fractions were dried and normalized to 0.2 μg/μL in 2%ACN in 0.1% TFA, and 5 μL per fraction were subject to no-FAIMS DDA analysis.

### Gas Phase Fractionation

For FAIMS-based gas phase fraction (GPF), 6 injections of tryptic peptides of each cell line were acquired by either DDA or DIA using the above settings. FAIMS user-set gas flow was set to 0 L/min, cooling gas flow was 5.0 L/min, and the temperature of both inner and outer electrodes 1 and 2 were set to 100 °C. We analyzed 1 μg of peptide with a range of serial CVs (−25V, −35V, −45V, −55V, −65V, −75V) using either DDA or DIA. For the segmented GPF, 6 fraction runs were acquired using a segmented DIA method previously reported (Brian C. Searle, 2018). We injected 1 μg of whole-cell digest using the same LC and DIA setting described above with different DIA separation ranges (400-500 m/z, 500-600 m/z, 600-700 m/z, 700-800 m/z, 800-900 m/z, 900-1000 m/z), all with 4 m/z isolation windows.

### DDA- and DIA-based library construction

The DDA raw files of FAIMS-DDA GPF and LPF were searched by MaxQuant (v1.6.10.43) (Cox & Mann, 2008) using UniProt human proteome reference (v20191105), which includes 74,788 protein entries with isoforms. The mass tolerance for precursor ions and fragment ions was set to 4.5ppm and 0.05Da, respectively. Methionine oxidation and N-terminal acetylation were selected as variable modifications. Both false discovery rates (FDRs) at the peptide and protein levels were set to 0.01. To generate the DDA spectral libraries, the DDA MaxQuant msms .txt output files were imported to Spectronaut (v14), at which time the “Generate Spectral Library from MaxQuant” function was used to create the spectral libraries with the same Uniprot reference (v20191105) as the FASTA file.

The DIA raw files of FAIMS-DIA GPF and LPF were searched by Spectronaut (SN). These searches were directly used by Pulsar (a search engine embedded in SN) to generate DIA-based spectral libraries via the “Generate Library from Pulsar / Search Archives” function. The default BGS factory settings, as well as the same Uniprot reference (v20191105) were used when generating all libraries.

### Database search for proteomics

We performed direct DIA database searches in three engines: Spectronaut (v14), EncyclopeDIA (v0.9.5) (Bekker-Jensen, Bernhardt, et al., 2020) and PEAKS studio (v10.6)(Ma et al., 2003). Trypsin and/or lysine C were selected as the digestion enzyme in all search engines. Carbamidomethylation was selected as a fixed modification, and methionine oxidation and N-terminal acetylation were selected as variable modifications. FDRs of PSMs and peptide/protein groups were set to 0.01. Fragmentation was set to HCD or CID/HCD. In Spectronaut, we chose MS2 as quantity MS-level, and data imputing was disabled. In PEAKS studio, we selected the sequence database search as direct DIA, with MS1 scans used for quantification. For EncyclopeDIA, we first converted the MS raw files to MZML files using MSConvertGUI (v.3.0.2) (Chambers et al., 2012). We used the Walnut GUI embedded in EncyclopeDIA as direct DIA. The normal target/decoy and overlapping DIA were selected, and ion charge range was set from 2 to 4.

### Interactive aligned time-based UMAP

Aligned-UMAP is used for dimensionality reduction of our temporal proteomic data. Briefly, MAPPER algorithm is used to create regularizer term, enforcing the constraint on how far related points can take different locations in embeddings at multiple time points. The data analysis pipeline was generated in Python v3.8 using open-source libraries (numpy, pandas, plotly, umap). For evaluating the software output and reproducibility, a demo of the Aligned-UMAP visualization is available at https://share.streamlit.io/anant-dadu/alignedumap-biomedicaldata.

### Single-cell RNA-Seq library preparation and sequencing

The KOLF2.1J iPSCs and differentiated iNeurons (day 28) were washed once with 1x PBS (Lonza) before adding TrypLE (Gibco) containing 10 units/ml of papain (Worthington). Cells were incubated at 37°C until cells appeared to physically detach from each other when viewed under a phase contrast microscope. The papain solution was aspirated, and cells were dissociated with neural maturation media supplemented with 10 μM Rock inhibitor, 60 units of DNase I (Worthington). The cell suspension was centrifuged at 160 g for 5 minutes. The cells were resuspended and washed three additional times in 1× PBS-0.04% BSA (Jackson Immunoresearch). Single-cell pellets were resuspended in 1× PBS-0.04% BSA, counted using an automated cell counter (Countess II), and the concentration was adjusted to 1×10^6^ cells/mL. Single cell RNA sequencing library preparations were performed using a 10X Genomics Chromium Single Cell 3’ Reagent kit V3.1 (PN-1000128) with 25,000 cells per condition loaded into separate lanes of a 10X Genomics chip G. The indexed libraries were combined and sequenced on an Illumina HiSeq2500. The data were processed through the 10X Genomics Cell Ranger 5.0.0 pipeline.

### Single-cell RNA-Seq data analysis

The sequencing count files from the 10x Genomics Cell Ranger pipeline were further subjected to data pre-processing, dimensionality reduction, clustering, and annotation using the python package Scanpy (v1.8) (Wolf et al., 2018). Count matrices were converted to the AnnData objects. Cells with < 300 genes detected, > 9000 total counts, > 10 % mitochondrial genes were removed. The data were logarithmically transformed and normalized to 10000 reads per cell. Highly variable genes were extracted, then the effects of total counts and the percentage of mitochondrial genes were regressed out and scaled to unit variance. Dimensionality reduction was performed by running principal component analysis. Samples from different batches were integrated into one single AnnData object. The batch effect was corrected using batch balanced KNN (v1.5.1)(Polanski et al., 2020). Cells were clustered using the Leiden graph-clustering method, which detects communities based on optimizing modularity (Traag et al., 2019). Clustering was iterated at the resolution ranging from 0.4 to 1.2. Uniform Manifold Approximation and Projection (UMAP) was used for visualization. The two-dimensional embedding of UMAP was calculated from the first 50 principal components with the following parameters: spread = 1 and minimum distance = 0.7. The cell identity of each cluster was annotated by calculating the overlap score between data-driven cluster marker genes and the reference marker gene list from PanglaoDB (Franzen et al., 2019) (**Supplemental Table S1**) followed by manual curation. The function of scanpy.tl.score genes was performed to calculate the average expression of a group of genes of interest.

## Results

### Comparison of library-based and direct-DIA in single-shot proteomics

In attempts to maximize proteome coverage, we first systematically evaluated singleshot DDA and DIA using spectral and sequence libraries generated by offline liquid phase fractionation (LPF) or gas phase fractionation (GPF), including both segmented and FAIMS-based GPF. To assess the proteome coverage in distinct cell types, we performed parallel experiments with peptides from both HeLa protein digest and processed lysates of two iPSC lines, WTC11 and KOLF2.1J. The tryptic peptides of the iPSCs were generated using an fully automated SP3 sample preparation method, followed by different fractionation techniques (i.e., FAIMS-GPF, offline LPF, segmented GPF, and GPF, as described in Searle et al. 2018 (Searle et al., 2018)) and data acquisition (i.e., DDA and DIA) strategies **(Fig. 1A)**. We didn’t observe a significant improvement when using offline LPF compared to FAIMS-based GPF, and discovered that the FAIMS-DDA GPF-based library resulted in more identified precursors than the FAIMS-DIA GPF-based library in both iPSC lines **(Supplemental Fig. S1C)**. Next, we sought to evaluate total peptide and protein group identifications using library-based DIA and direct-DIA (dDIA). We performed single-shot proteomics without using FAIMS of each cell line with either DIA or DDA acquisition, followed by database searches using the previously generated libraries (library-DIA) or searched concurrently with the fractionation raw files (dDIA). These results suggested that (1) DIA with or without fractionated library consistently identified more peptides and proteins than DDA in single-shot proteomics; (2) the use of an in-house segmented GPF library yields higher identification than that generated through Searle et al GPF, suggesting that an instrument- and method-specific library is crucial for DIA; (3) FAIMS-DIA GPF identified more peptides and proteins than other libraries; (4) dDIA displayed better proteome coverage than library-DIA in all three cell lines **(Fig. 1B)**. When comparing all peptides and proteins identified in dDIA and library-DIA, more peptides and proteins were exclusively identified by dDIA than library-DIA. For example, 31.8% of total protein groups were only identified by dDIA in WTC11 cells while 3.2% of those were only found using library-DIA **(Fig. 1C)**. Taken together, our findings suggest that FAIMS-DIA GPF combined with dDIA provides the deepest proteome coverage in single-shot proteomics.

**Fig.1.**
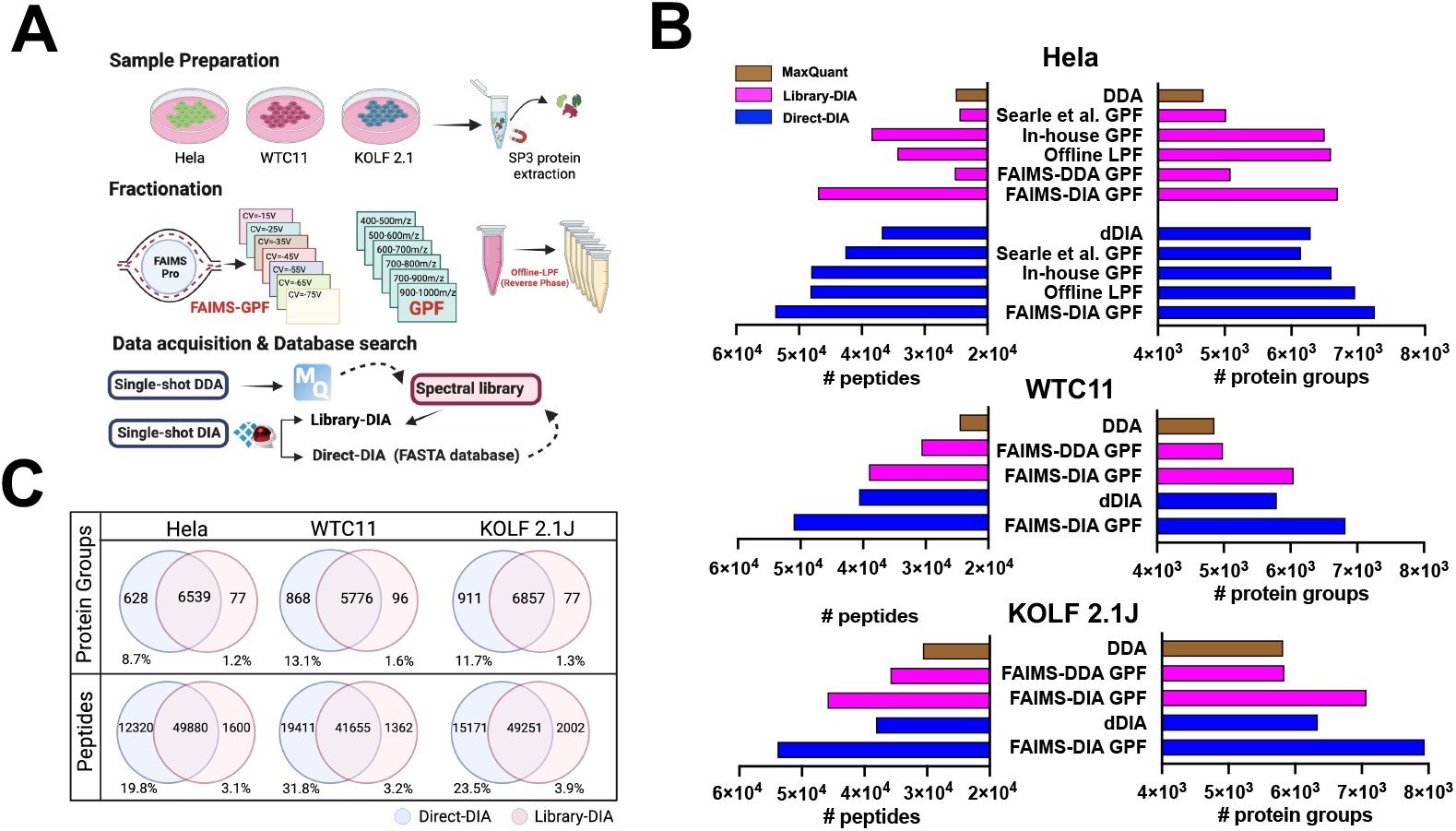
Comparison of library-based and direct-DIA in single-shot proteomics. **(A)** Schematic diagram of sample preparation, fractionation, library generation, and database search strategy. (**B)** Bar chart shows the number of peptides and protein groups identified in single-shot DDA and DIA runs in Hela cell digest, WTC11, and KOLF2.1J iPSCs using MaxQuant-based DDA, library-DIA, and direct-DIA. **(C)** Venn diagrams display the number and percentage of protein groups or peptides exclusively detected in library-DIA or direct-DIA.

### Comparison of FAIMS-DIA with a range of single CVs

Using FAIMS, it is possible to modify compensation voltage (CV), determining the subset of ions which pass through the device, and ultimately downstream detection of peptides. To determine which CV led to the largest number of identified peptides and protein groups, we compared FAIMS-DIA with single CVs from −15V to −85V with 10V steps and DIA without FAIMS. Both methods used a 2-hour LC gradient, with MS runs searched against a database individually via dDIA. Strikingly, we identified more protein groups in both individual −35V and −45V runs compared to DIA without FAIMS **(Fig. 2A)**, despite FAIMS-DIA identifying fewer peptides than DIA without FAIMS at almost every CV tested **(Fig. 2B)**. When comparing the charge of peptides identified at different CVs, we discovered that CV = −25V, −35V and −45V favor 2+ ions, while CV = −55V, −65V and −75V favor 3+ ions. The dominant ions for DIA without FAIMS were 2+ **(Fig. 2C)**. Out of the combined pool of peptides detected by each CV of FAIMS-DIA, there were 10792 (18.5%) and 7221 (12.4%) peptides exclusively identified at −35V and −45V of FAIMS-DIA, respectively **(Fig. 2D and Supplementary Fig. S2A)**. Similarly, we observed that 11774 (22.6%) and 7726 (14.8%) peptides were exclusively identified at −35V and −45V in FAIMS-DDA, respectively **(Supplementary Fig. S2B-C)**.

**Fig.2.**
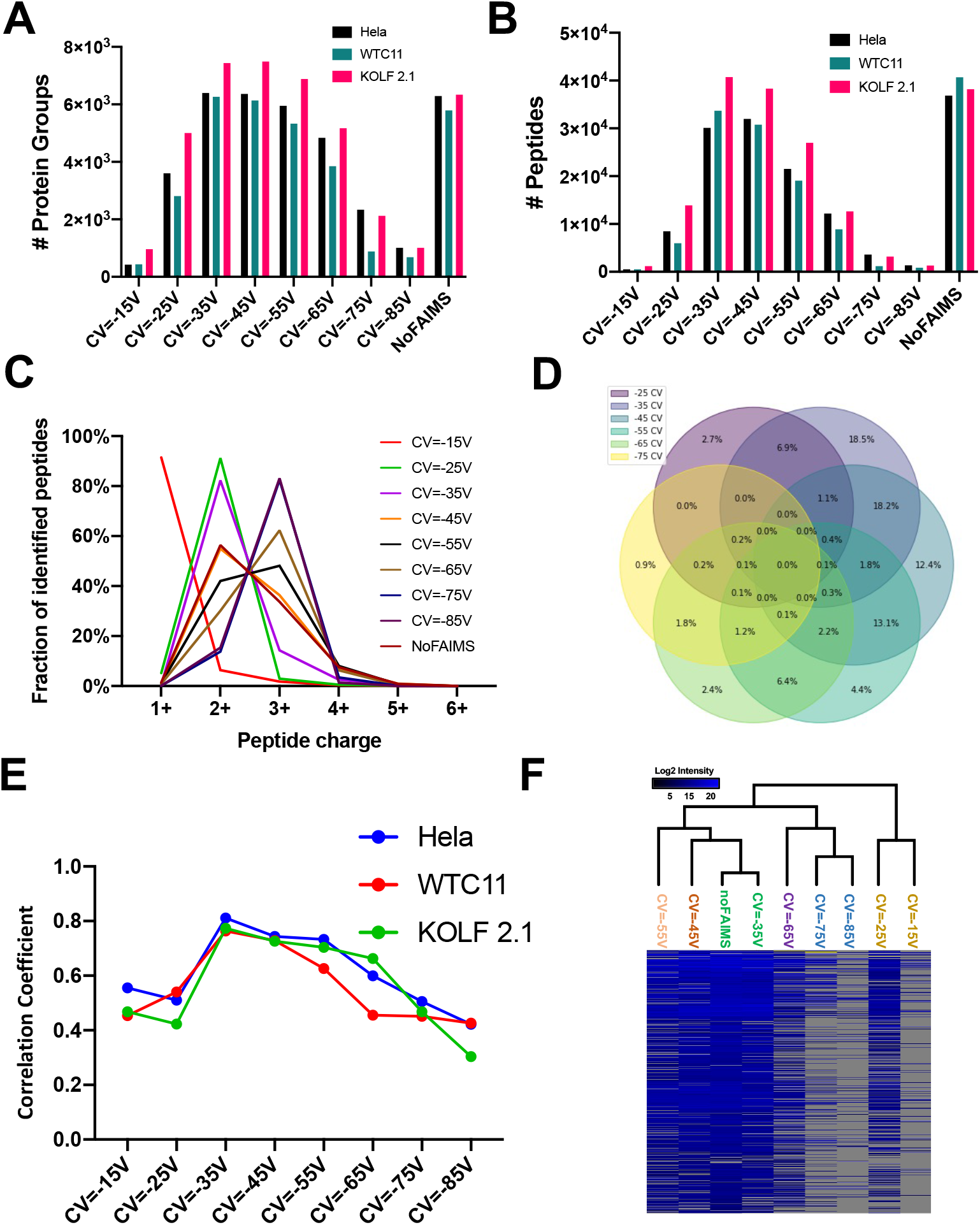
Comparison of FAIMS-DIA with a range of single CVs. **(A-B)** Bar charts shows the number of protein groups **(A)** and peptides **(B)** identified using 8 FAIMS single-CV-DIA (−15V to −85V, 10V stepping) and noFAIMS-DIA approaches in Hela digest, WTC11, and KOLF 2.1J iPSCs. **(C)** Fraction of identified peptides with a range of charges (1+ to 6+) by 8 FAIMS single-CV-DIA and noFAIMS-DIA in Hela digest. **(D)** Venn diagram of the percentage of peptides exclusively identified in 6 FAIMS single-CV-DIA runs (−25V to −75V). **(E)** Pearson correlation coefficients of protein abundance detected by noFAIMS-DIA and 8 FAIMS single-CV-DIA (−15V to −85V) in Hela digest, WTC11 and KOLF2.1J iPSCs. **(F)** Dendrogram cluster of proteins identified by noFAIMS-DIA and 8 FAIMS single-CV-DIA (−15V to −85V) in KOLF 2.1 iPSCs.

To evaluate the consistency of protein abundance quantified across conditions, we performed a linear regression of log2-transformed intensities of proteins identified in each tested CV against DIA without FAIMS. Runs with CV = −35V (FAIMS (−35V)-DIA) showed the highest correlation coefficient with DIA without FAIMS in all three cell lines **(Fig. 2E)**. Similarly, unsupervised dendrogram cluster analysis confirmed that the protein abundance quantified at CV = −35V had the greatest similarity with those acquired using DIA without FAIMS in KOLF2.1J iPSCs **(Fig. 2F)**. In summary, our results suggest that FAIMS (−35V)-DIA could be an alternative approach to achieve similar proteome coverage compared to DIA without FAIMS, even without the use of large, sample-specific libraries.

### Comparison of database search engines for direct-DIA

Since we have demonstrated that dDIA yields more identifications than library-DIA, we sought to explore three commonly used database search engines supporting dDIA: Spectronaut (SN), Walnut-based EncyclopeDIA (EN) and PEAKS studio. One of the benefits of single-CV FAIMS is that nearly all database search algorithms can process raw files generated from runs using a single applied CV; however, MS runs with multiple CVs must be deconvoluted and converted into a single file prior to database search (Hebert et al., 2018). Thus, we tested the overall search engine performance using KOLF2.1J iPSC single-shot whole-proteome data acquired by FAIMS (−35V)-DIA (*n* = 6). SN identified and quantified the most protein group IDs, with less than 30% missing values **(Fig. 3A)**. Additionally, SN identified nearly 60,000 peptides, considerably outperforming both EN and PEAKS **(Fig. 3B)**. Venn diagrams of protein and peptide detection between search engines confirmed that SN provided the highest coverage and most exclusive peptides among all tested algorithms **(Fig. 3C-D)**. When comparing the protein quantitation readouts by different search engines, SN and PEAKS showed the highest correlation (mean r^2^ = 0.85), while SN and EN had the lowest correlation (mean r^2^ = 0.78) **(Supplemental Fig. S3 A-C)**.

**Fig.3.**
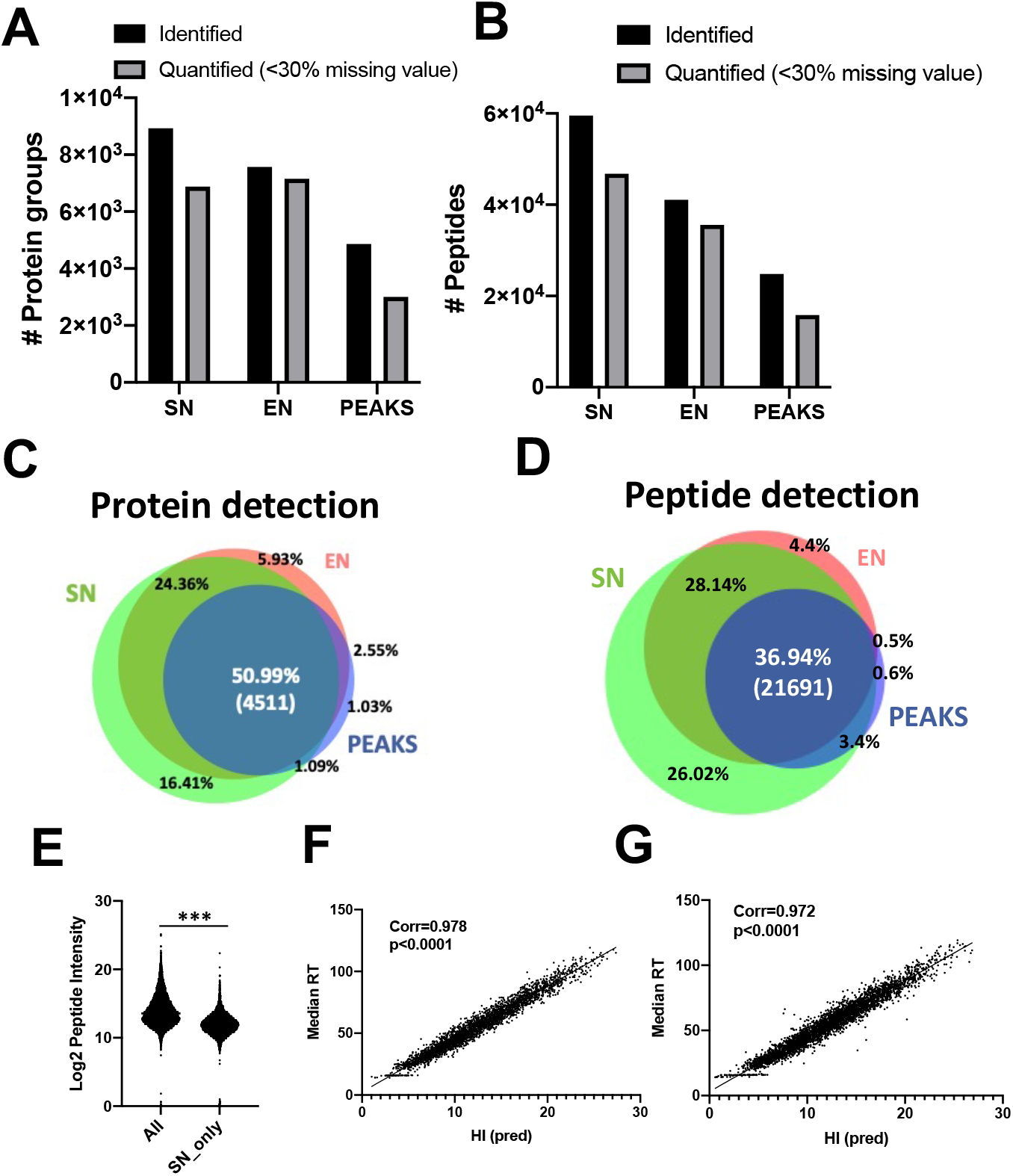
Comparison of database search engines for direct-DIA. **A-B)** Bar charts show the number of identified and quantified protein groups **(A)** and peptides **(B)** in KOLF2.1J iPSCs (*n* = 6) using SN, EN and PEAKS. **(C-D)** Venn diagrams of the overlapping total identified protein groups **(C)** and peptides **(D)** using SN, EN and PEAKS. **(E)** Log2 transformed intensity of all peptides identified by SN and the peptides exclusively identified by SN. **F-G)** Correlation of median retention time across 6 biological replicates and predicted hydrophobicity index of all peptides identified by SN **(F)** and the peptides exclusively identified by SN **(G)**.

Next, we sought to ensure that the large number of peptides exclusively detected by SN were not the result of false-positive identification. We observed relatively low intensity of the peptides exclusively identified in SN compared to commonly identified peptides **(Fig. 3E)**. Despite the low abundance of these peptides, the median retention time and predicted hydrophobicity (HI) of all peptides or SN-exclusive peptides correlated with Pearson’s coefficients of 0.978 and 0.972, respectively **(Fig. 3F-G)**. Further, MS/MS transition spectra of select peptides, uniquely identified by SN, confirmed multiple true fragment ions from each peptide **(Supplemental Fig. S3D-H)**. Taken together, this data demonstrates that dDIA searches in SN indeed result in the best proteome coverage.

### Temporal proteomic profiling of iPSC-derived neurons

To further evaluate our method of FAIMS-DIA in conjunction with library or dDIA data analysis, we profiled the proteome of iPSC-derived neurons at 7 time points over the course of a 28-day differentiation period (day0 [d0], d3, d7, d10, d14, d21, d28). We adopted the neurogenin-2 (NGN2) transcription factor-based neuronal differentiation protocol, in which iPSCs are transduced with doxycycline (DOX)-inducible NGN2. These iPSCs can then be differentiated into neuron-like cells by addition of doxycycline to the cell culture media (Fernandopulle et al., 2018; Zhang et al., 2013). We included 6 biological replicates per time point during neuron differentiation, and data was acquired by the above DIA approaches, FAIMS (−35V)-DIA and DIA without FAIMS. The MS raw files of FAIMS (−35V)-DIA were database-searched by dDIA in SN while the DIA without FAIMS raw data was searched with or without the previously generated FAIMS-DIA GPF library (Schematic shown in **Fig. 4A**). To validate the neuron differentiation protocol, we performed live cell analysis of neurite morphology and outgrowth using Incucyte. We observed neuron-like cells at d3, synaptic formation at d14, mature neuronal morphology at d21, and peak neurite growth occurring at d25 **(Fig 4B and supplemental Fig. S4A)**.

**Fig 4.**
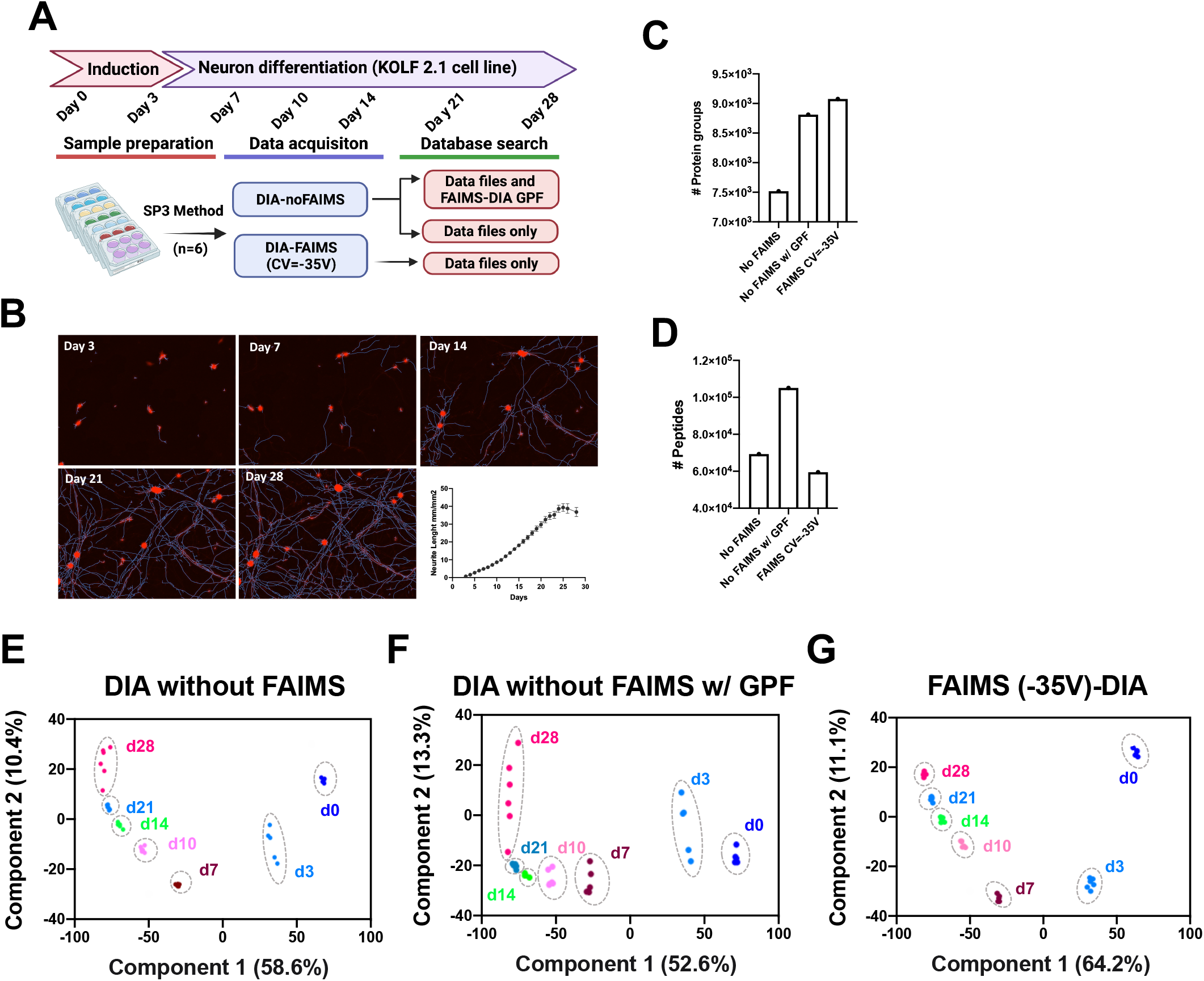
Temporal proteomic profiling of iPSC-derived neurons. **(A)** Schematic diagram of a 28-day neuron differentiation of KOLF2.1J-derived i Neuron. Six biological replicates (*n* = 6) of each time point were included and subjected to noFAIMS-DIA and FAIMS (−35V)-DIA proteomic analysis. Database search was conducted with or without FAIMS-DIA GPF. **(B)** Representative live cell images of neurite outgrowth on d3, d7, d14, d21, d28. Scatter plot shows the neurite length during the 28-day neuron differentiation. C-D) Bar charts show total number of protein groups (E) and peptides (D) identified by different direct-DIA approaches: noFAIMS, noFAIMS with GPF, FAIMS CV=-35V. **(E-G)** Principal component analysis (PCA) of the KOLF2.1J-derived neurons across 7 time points acquired by noFAIMS **(E)**, noFAIMS with GPF **(F)**, FAIMS CV=-35V **(G)**.

Proteomic analysis of differentiating neurons indicated that, although DIA without FAIMS with GPF library identifies a greater number of peptides, FAIMS (−35V)-DIA identifies more protein groups **(Fig. 4C-D)**. Reproducibility was assessed by calculating the coefficient of variation percent (CV%) among replicates. The CV% of FAIMS (−35V)-DIA was significantly lower than the other two approaches **(Supplemental Fig. S4B)**. Principal component analysis (PCA) revealed tighter clustering of replicates collected at different time points when using FAIMS (−35V)-DIA. Interestingly, we found that the use of a GPF library resulted in greater inter-sample variation, compared to DIA without FAIMS datasets searched with or without spectral library, based on PCA clustering **(Fig. 4E-G)**. Although the FAIMS-based approach increased the total identified proteins, the question of whether this method reflects dynamic changes in protein abundance remains unclear. We selected panels of mature neuronal, synaptic, and neural stem markers to visualize differential expression throughout the longitudinal profiling, observing similar pattern between FAIMS-based DIA and regular DIA without FAIMS, with strikingly fewer missing values detected when using our FAIMS-based method **(Supplemental Fig. S4C)**. Thus, FAIMS (−35V)-DIA outperformed the other two approaches in both the larger number of proteins identified and superior reproducibility.

Next, we used the FAIMS (−35V)-DIA datasets to assess the efficiency of neuronal differentiation efficacy in KOLF2.1J-derived Neurons **(Supplemental Table S2)**. As expected, protein expression of pluripotency and neural precursor markers POU5F1 (OCT4) (Pesce & Scholer, 2000), PODXL (TRA-1-60)(Draper et al., 2002; Henderson et al., 2002), and SOX2 (Botquin et al., 1998; Ellis et al., 2004) were significantly reduced at the early stage of neuron differentiation **(Fig. 5A)**. The mature neuron markers, MAPT (Tau) (Binder et al., 1985) and NEFM (Julien & Mushynski, 1998) were rapidly upregulated from d0 to d7, plateauing later in neuronal differentiation (d14-d28) **(Fig. 5B)**. Rapid assembly of synaptic machinery was demonstrated by steadily upregulated synapse markers, including SYN1 (De Camilli et al., 1983), SNAP25 (Hess et al., 1992), and SYT1 (Fernandez-Chacon et al., 2001) (**Fig. 5C)**. Most importantly, our results indicate the presence of cortical-like neurons; protein expression of upper cortex layer markers (e.g., RELN, CUX1, CACNA1E) and deeper cortex layer markers (e.g., NECAB1, NTNG2) were enhanced over the course of differentiation **(Fig. 5D)** (Zeng et al., 2012). Differential expression analysis of iPSCs (d0) and mature neurons (d28) revealed that more than 65% of total proteins were significantly altered (Adj-p<0.05 and >1.5-fold difference between groups), including upregulated MAP2, MAPT, CTNT1, and downregulated MCM5, LIN28A, APOE **(Fig. 5E)**. Gene ontology (GO) KEGG pathway analyses of all upregulated proteins identified neurotransmitter secretion, axon guidance, and synapse signaling pathways, whereas analyses of downregulated proteins identified DNA replication, rRNA processing, and cell cycle signaling cascades **(Fig. 5F-G)**. Furthermore, we performed integrated pathway analysis considering the fold-change of all proteins and discovered activated synaptic and axon, mTOR, AMPK, and PKA signaling pathways (positive Z-score) and inhibited EIF2, chromosomal replication, and PTEN signaling (negative Z-score) **(Fig. 5H)**. Strikingly, the synaptogenesis signaling pathway was the most significantly enriched pathway **(Supplemental Fig. S5A)**. Moreover, we visualized the protein expression trajectory of select ADRD genes reported in the iNDI project during neuron differentiation (Ramos et al., 2021) **(Supplemental Table S3)**. Protein expression of Alzheimer’s Disease-related genes (e.g., *ABCA7, APOE*), Parkinson’s Disease-related genes (e.g., *DNAJC6, DNMT1*), and frontotemporal dementia/amyotrophic lateral sclerosis-related genes (e.g., *ANXA11, CHMP2B*) were found to be altered with neuronal differentiation **(Fig. 5I)**. Addtionally, for data visuliaztion and browsing, we made a interactive 3D plots of aligned UMAP based on the longitudinal proteomic datasets (d0-d28), including different sub-feature sets of specfic markers for mature neurons, synapase, cholinergic and glutamatergic neurons (https://share.streamlit.io/anant-droid/singlecellumap) **(Supplemental Fig. S5B)**. In summary, our in-depth proteomic profiling suggests that NGN2-based differentiation of KOLF2.1J iPSCs results in highly differentiated, cortical-like neurons.

**Fig.5.**
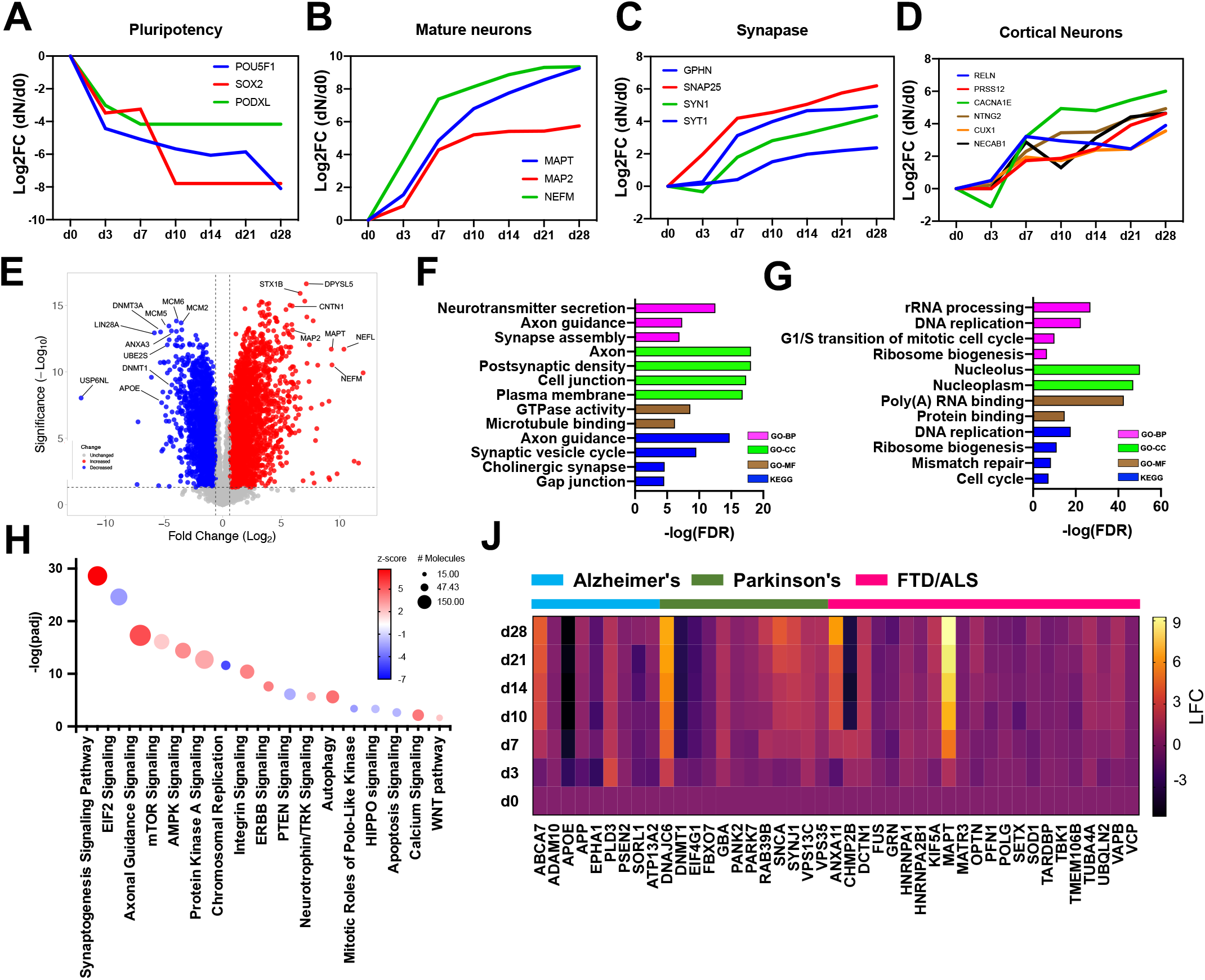
Functional annotation and pathway analysis of differentially expressed proteins during neuron differentiation. **(A-D)** Protein expression trajectory of select pluripotency markers **(A)**, mature neuron markers **(B)**, synapse markers **(C)**, cortical neuron markers **(D)** from d0 to d28. **(E)** Volcano plot shows the differentially expressed proteins comparing d28 neurons to iPSCs (d0); top 10 most significantly altered proteins were labeled; up-regulated proteins (FC > 2, padj<0.05) are color-coded in red, and down-regulated proteins (FC < 0.5, padj<0.05) are color-coded in blue. **(F-G)** Gene Ontology (GO) and KEGG pathway analysis of up-regulated proteins **(F)** and down-regulated proteins **(G)**. **(H)** Bubble chart incorporating the integrated pathway analysis of all proteins; Z-score was color-scaled (positive score indicates activated pathways while negative score indicates inhibited pathways); the bubble size indicates the number of proteins involved in respective signaling pathway. **(J)** Heatmap shows the log2-transformed fold changes (relative protein expression against d0) of select ADRD genes over the course of neuron differentiation.

### Single-cell RNA sequencing and cluster analysis of iPSC-derived neurons

To validate our observations from MS, we performed single-cell RNA sequencing (scRNA-seq) of d0 iPSCs and d28 iNeurons. This yielded an average of 6839 cells per sample with an average of 2805 median genes per cell. As expected, iPSCs and neurons show distinct clustering in two-dimensional UMAP space **(Fig. 6A)**. More importantly, the majority of the iNeurons appear to be mature, with comparatively small fractions of neural precursors and stem cells **(Fig. 6B)**. Further, we used a fuzzy K-mean clustering algorithm to the group of proteins identified by MS into 50 distinct clusters based on their protein expression trajectories, reflected by the relative foldchange (FC) of each time point compared to d0 iPSCs. Then, 50 clusters were subdivided into “up” (11 clusters), “down” (11 clusters), and “not-significant (NS)” (28 clusters), depending on their differential expression in d28 neurons compared to iPSCs. Pathway analyses of these clusters revealed that “up” clusters contained cortical and glutamatergic neuron markers involved in synapse and neurogenesis, whereas “down” clusters contained proteins involved in DNA replication and cell division. Interestingly, expression of proteins in proteasome, mitochondria, and mRNA processing largely remains unchanged, falling into the “NS” cluster **(Table 1, supplemental fig. S6)**. Clusters 40 and 48 contain two neuronal markers, MAPT and MAP2, respectively. We confirmed their gene expressions were highly enriched in iNeurons compared to iPSCs using scRNA-seq; strikingly, module expression of all genes identified clusters 40 and 48 were well-correlated with the gene expression of *MAPT* and *MAP2*, respectively **(Fig. 6C-D)**. In contrast, neural precursor markers, *POU5F1 and SOX2* (belonging to cluster 15 and 25, respectively), were highly expressed in iPSCs; module expression of both clusters showed similar patterns **(Fig. 6E-F)**. Given the concordance between both proteomic and scRNAseq-level gene expression analysis, our results further support the validity of our longitudinal proteomics in neuron differentiation.

**Fig.6.**
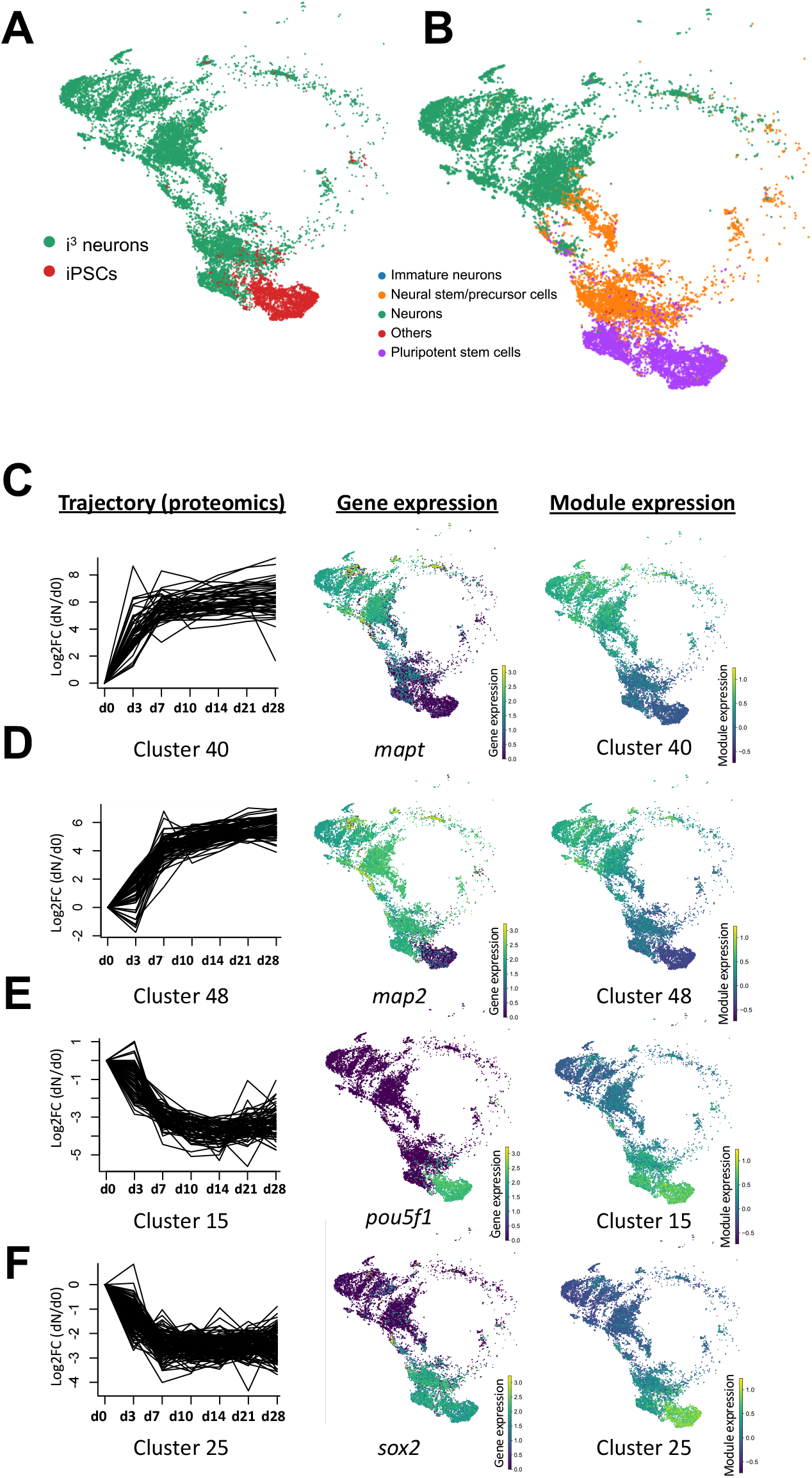
Single-cell transcriptomic analysis of iPSC-derived neurons. **(A)** Uniform manifold approximation and projection (UMAP) analysis of KOLF 2.1 iPSCs (d0) and derived d28 neurons. **(B)** UMAP analysis shows annotated clusters indicating neurons (green), immature neurons (blue), neural precursors (orange), stem cells (purple). **(C-F)** Representative examples of K-mean cluster analysis based on the protein expression trajectories over the course of neuron differentiation (left panel); UMAP plot of select genes in respective clusters (center panel); UMAP plot of module expression score of respective clusters (right panel). Four select clusters (2 up-regulated and 2 down-regulated) and 4 markers (one for each cluster) were shown including cluster 40 (representative marker: *MAPT*) **(C)**, cluster 48 (representative marker: *MAP2*) **(D)**, cluster 15(representative marker: *POU5F1*) **(E)**, cluster 25(representative marker: *SOX2*) **(F)**.

**Table 1.**
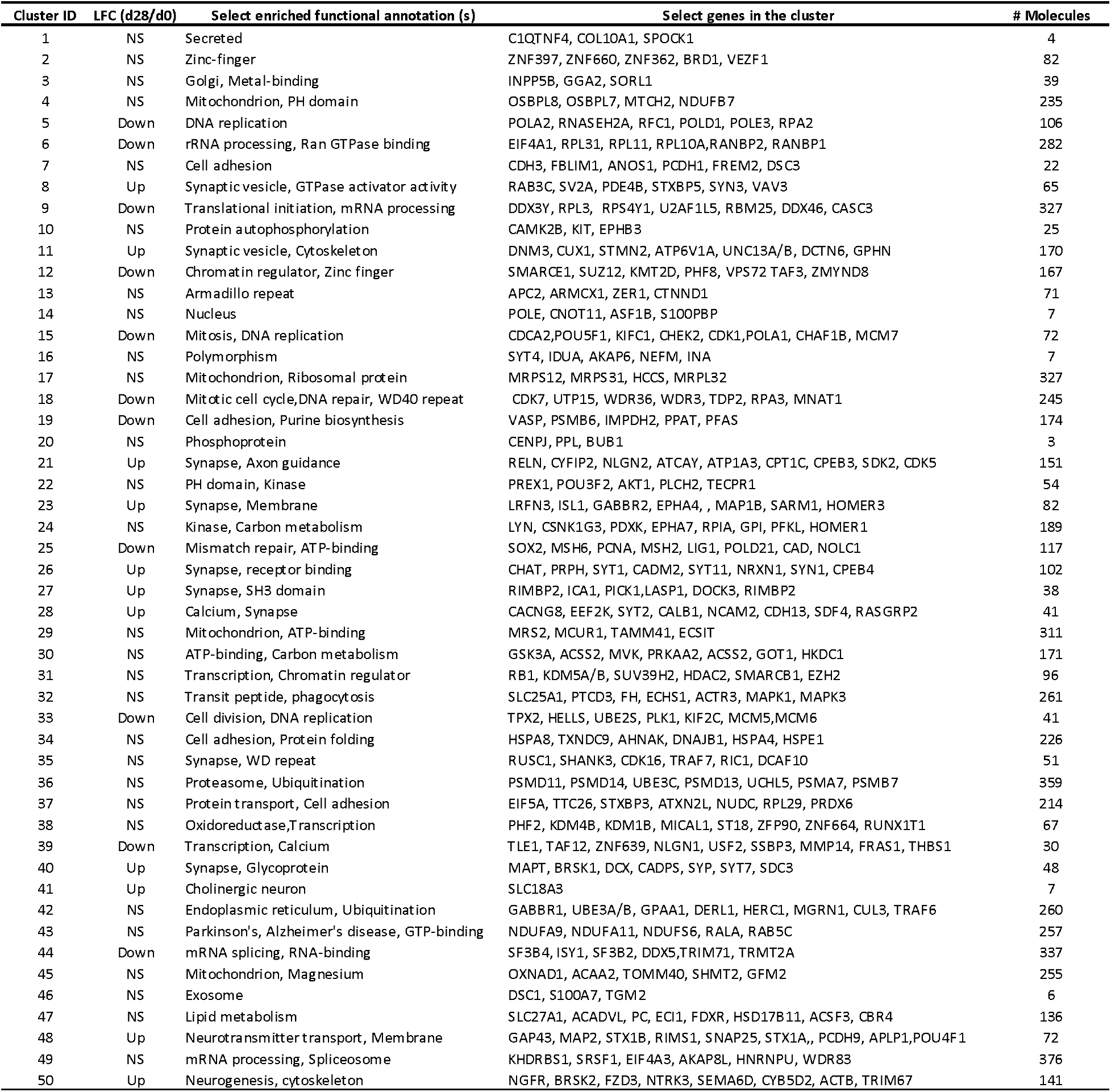
Cluster analysis reveals protein expression dynamics and enriched pathway during neuron differentiation.

Next, we applied our protein clusters to identify potentially novel genes that are related to neuron maturation. For example, we selected 31 representative proteins from 6 “up” clusters (Draper et al., 2002; Frade & Ovejero-Benito, 2015; Pino et al., 2020; Traag et al., 2019; Zhang et al., 2013), of which 6 have been reported as neural markers, 18 of the remaining 25 proteins were found to be specifically expressed in the brain based on the tissue atlas database (https://www.proteinatlas.org/) (**Fig. 7A**). This included proteins which have known neuronal roles, but previously uncharacterized expression specificity to mature neurons (e.g. LSAMP, CTNNA2). We also identified proteins involved in putatively ubiquitous processes such as RNA metabolism (e.g., ELAVL2 (Berto et al., 2016)) and organelle dynamics (e.g., GDAP1 (Huber et al., 2013) and RTN1 (Gong et al., 2017)), revealing that these cellular phenomena are important to both neurodevelopment and mature neuron function. Additionally, single cell-transcriptome analysis revealed that, compared to iPSCs, the majority of d28 neurons expressed relatively high levels of *ELAVL2, GDAP1*, and *RTN1*, and the cortical neuron marker *CUX1*, which showed similar gene expression patterns (**Fig. 7B**). Taken together, we find that scRNA-seq combined with cluster analysis of protein expression data may unveil potentially novel correlates of neuron differentiation.

**Fig.7.**
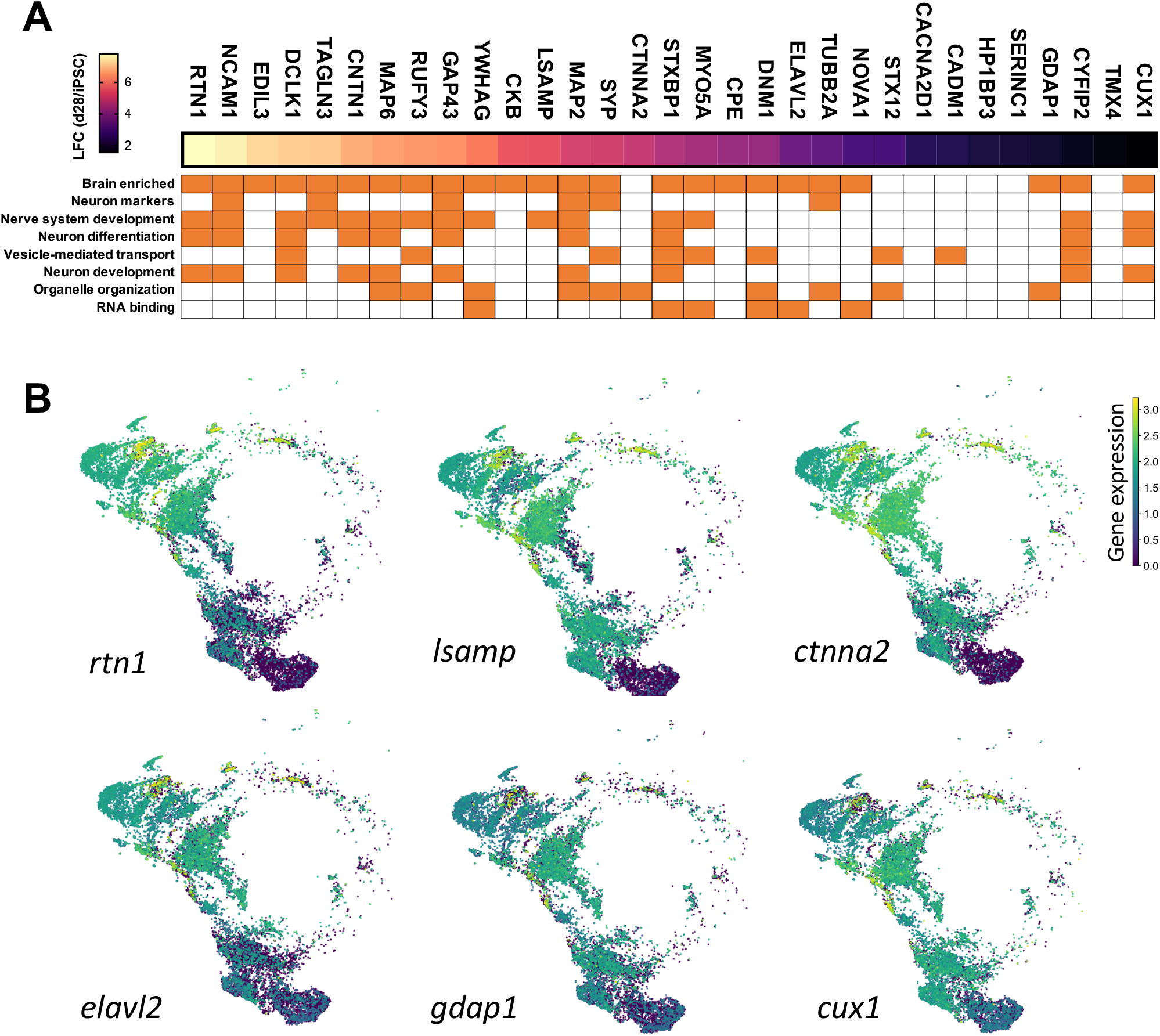
Potentially novel neuron differentiation-related proteins. **(A)** 31 select genes from 5 up-regulated clusters were annotated by tissue atlas database and GO term and classified into different categories. Heatmap indicating the log2 transformed fold change of given protein expressed in d28 neurons compared to iPSCs. **(B)** scRNAs-seq UMAP plots of representative genes not previously associated with neuron differentiation, such as *RTN1, LSAMP, CTNNA2, ELAVL2, GDAP1, CUX1* (a cortical neuron marker), indicating they are highly expressed in iNeurons.

## Discussion

In this study, we systematically investigated different single-shot proteomics strategies to characterize thousands of proteins accurately and reproducibly in a high-throughput fashion using a fully-automated workflow. We found that, of the approaches we evaluated, single CV = −35V FAIMS combined with a direct DIA database search strategy provided the deepest proteome coverage and highest reproducibility. Further, we applied this approach to map the proteomic signatures of differentiating KOLF2.1J iPSC-derived neurons over time and demonstrated that mature and cortical neuron markers are highly expressed at the protein level in d28 neurons. Leveraging scRNA-seq, we validated that the majority of cells at d28 are indeed mature neurons, and the differential expression of many neural and stem cell markers were also observed at the transcript level.

The two major data acquisition approaches for shotgun proteomics are DDA and DIA. Our results suggest all DIA methods (e.g., library or direct DIA, with or without FAIMS) outperform DDA methods, aligning with many previous reports (Dorte B. Bekker-Jensen, 2020; Searle et al., 2018). Conventional DIA utilizes DDA-based spectral libraries, with larger libraries often increasing the total identification (Pfammatter et al., 2021). Recently, DIA-based chromatogram libraries have been proven to increase identification, reduce missing values, and have better reproducibility over sample-specific DDA libraries (Pino et al., 2020; Searle et al., 2018; Searle et al., 2020). Our study demonstrates that direct-DIA outperforms library-based approaches, which usually require pooled samples from all specimens to build libraries. This finding is especially relevant for large-scale initiatives such as iNDI, in which samples are generated continuously over protracted periods of times, precluding a library-based approach. Strikingly, we showed inter-sample variation dramatically amplified when adding the GPF library **(Fig. 4E, F)**. Although previous studies have demonstrated that multiple-CV FAIMS returns more identifications than single CV-FAIMS in DDA proteomics (Hebert et al., 2018; Muehlbauer et al., 2020), multiple-CV FAIMS does not significantly improve proteome coverage compared to single-CV FAIMS when using DIA (Bekker-Jensen, Martinez-Val, et al., 2020). This is possibly because DIA requires higher quality MS2 spectra than DDA, and cycle time is often split between the multiple CVs during MS2 acquisition. Another advantage of single-CV FAIMS DIA is that all engines can process single-CV files for database searches, while some search engines are incompatible with multiple-CV FAIMS data. Despite the inherent bias of peptide sampling in single-CV FAIMS compared to multiple-CVs and without using FAIMS, we show that the overall protein quantification of optimized single-CV (−35V) correlates strongly with without using FAIMS which generally is recognized as the ‘ground truth’. Direct-DIA is commonly based on pseudo MS2 spectrum alignment and deep learning, such as DIA-Umpire (Tsou et al., 2015), DIA-NN (Demichev et al., 2020), EN (Searle et al., 2018), SN, PEAKS; therefore, MS1 scans are required for the direct-DIA approach. In addition to being the two best-performing algorithms, SN and PEAKS offer the advantage of being recent, commercial software with ongoing technical support and version updates. SN identifies the greatest proportion of unique peptides that are not detected by other search engines.

There are other data acquisition factors and LC settings we did not attempt to optimize in our current study. For example, we used a normal range of 400-1000 *m/z* with 8 m/z isolation windows for DIA, as prior data have suggested that acquisition settings such as range and window, as well as LC parameters including flow rate, do not significantly affect proteome depth (Kawashima et al., 2019). To maximize total identification within a reasonable timeframe, we adopted conventionally used parameters in studies evaluating FAIMS, suggesting 1 μg peptide injection with a 2-h LC gradient (Hebert et al., 2018). Exploring the effect of lower peptide injection and shorter gradient on total identifications may be helpful to evaluate in the future. We have tested this approach on two Orbitrap Eclipse instruments with more than 2000 MS runs and have observed consistent and reproducible results. FAIMS also provides several non-analytical advantages, including a reduction in instrument cleaning cycles. We acquired upwards of 100 samples without detecting significant single suppression, whereas instruments without FAIMS need to be cleaned every 30-50 samples (Poulos et al., 2020). Although this FAIMS-assisted DIA approach has been validated for the iNDI project, it has great potential to be applied to any large-scale study seeking reliable, reproducible, and accurate proteomics-based quantitation.

During the 28-day neuron differentiation, our live cell images of fully differentiated neurons (d25-d28) suggest mature neurite morphology and complete outgrowth. The longitudinal protein expression patterns of KOLF2.1J-derived neurons are similar to previously reported induced cortical neurons (Burke et al., 2020; van de Leemput et al., 2014), supporting the maturation of our^-^neurons. Many studies rely on transcriptomic data, so our proteomic data is complementary to prior datasets and provides a rich resource for future studies. Strikingly, 65% of proteins are differentially expressed over the course of neuron differentiation and maturation. Although this is a considerably higher fraction than found in other reports, similar molecular functions of identified proteins have been observed (Lindhout et al., 2020). More importantly, a majority of ADRD genes are differentially expressed during differentiation, indicating that this cellular model may be appropriate for modeling the effects of genetic variation associated with neurodegenerative disorders (Ramos et al., 2021). For easy data access, we bulit a webapp for better visualization and data browsing of this dataset using time-based interactive 3D plots.

The cluster analysis of protein expression trajectories deconvolutes genes involved in different stages of neuron development, such as pluripotency, synaptic formation, and neuronal maturation. The pluripotency markers OCT4 and SOX2 were substantially inhibited as early as d7 and d10. Mature neuron markers (e.g. MAP2 and NEFM) peak at d10, whereas the expression of many synaptic and cortical markers (e.g. SYN1, SNAP25, CUX1, RELN, NTNG2) rapidly increased until d10 and continued to increase in expression throughout the course of the experiment to d28. Further, we plotted the temporal transcriptome datasets from CORTECON (van de Leemput et al., 2014), which showed that some of the cortical neuron markers (e.g. *RELN, CACNA1E*) steadily increased as far out as day 77. These results indicate that although cortical development cannot be fully recapitulated *in vitro*, our d28 neurons can still mimic general cortical neuron characteristics (**Supplemental Fig. S7**). By leveraging scRNA-seq data, we not only confirmed that a majority of differentiated neurons represent mature cortical neurons, but also validated our observations from the cluster data generated by our proteomics dataset. The high expression patterns of protein clusters related to neuron maturation in d28 neurons is supported by single-cell transcriptome module expression scores **(Fig. 6C-D)**, suggesting a high correlation between proteomics and scRNA-seq data. In summary, for the first time, by combining proteomic and scRNA-seq, we systematically characterized and validated the KOLF 2.1J-derived cortical-like neurons as an ideal model for functional studies of ADRD genes in the iNDI project.

## Data availability

The MS raw and peaks files of the whole-cell proteome have been deposited to ProteomeXchange *via* PRIDE, project accession ID: PXD029902. All raw single-cell RNA-seq results have been deposited in the National Center for Biotechnology Information Short Read Archive, SRA accession ID: SRP347436.

## Code availability

Our code for aligned UMAP of temporal proteomic data is publicly available at https://github.com/anant-droid/SingleCellUMAP to facilitate replication and future expansion of our work. The repository is well documented and includes a description of the data pre-processing, statistical, and machine learning analysis used in this study (https://umap-learn.readthedocs.io/en/latest/aligned_umap_basic_usage.html).

## Acknowledgment

We thank Drs. Avindra Nath and Jeffrey Kowalak for supervising and overseeing the proteomic center within National Institute of Neurological Disorders and Stroke (NINDS)/NIH. We thank Dr. Sarah Hill and Maia Parsadanian from NINDS/NIH for providing the WTC11 iPSC line. We thank Sahba Seddighi for providing the informatic analysis assistance. We thank Kailyn Anderson and Matthew Nelson for providing the tissue culture assistantce.

## Author contributions

Y.A.Q., M.E.W. and L.R. conceptualized the research study. L.R. and Y.A.Q. conducted the sample preparation, mass spectrometry-based proteomics analyses, database search and downstream informatic analyses. L.R., E.L., D. R., M.S. and M.F. performed the cell culture and iPSC-neuron differentiation. L.P., E.L., J.L. and S.L.C. performed the single-cell sample preparation and sequencing, and data analyses. L.P., A.D., F.F. and M.A.N. conducted the longitudinal biomedical datasets webapp and performed statistical analyses. J. S. managed experiment supplies and data storage. L.R., C.P., and Y.A.Q. drafted the manuscript. Y.A.Q., M.E.W., P.N., M.R.C. and A.B.S. supervised the project. All authors have read and agreed to the published version of the manuscript.

## Supplemental materials

**Supplemental Table S1**. Reference marker gene list used for cluster annotation in single-cell transcriptome.

**Supplemental Table S2**. Normalized protein intensity in temporal proteomic profiling of KOLF2.1J differentiated neurons.

**Supplemental Table S3**. Fold changes of protein expression of ADRD genes during a 28-day differentiation in KOLF2.1J-derived neurons.

**Supplemental Fig. S1.**
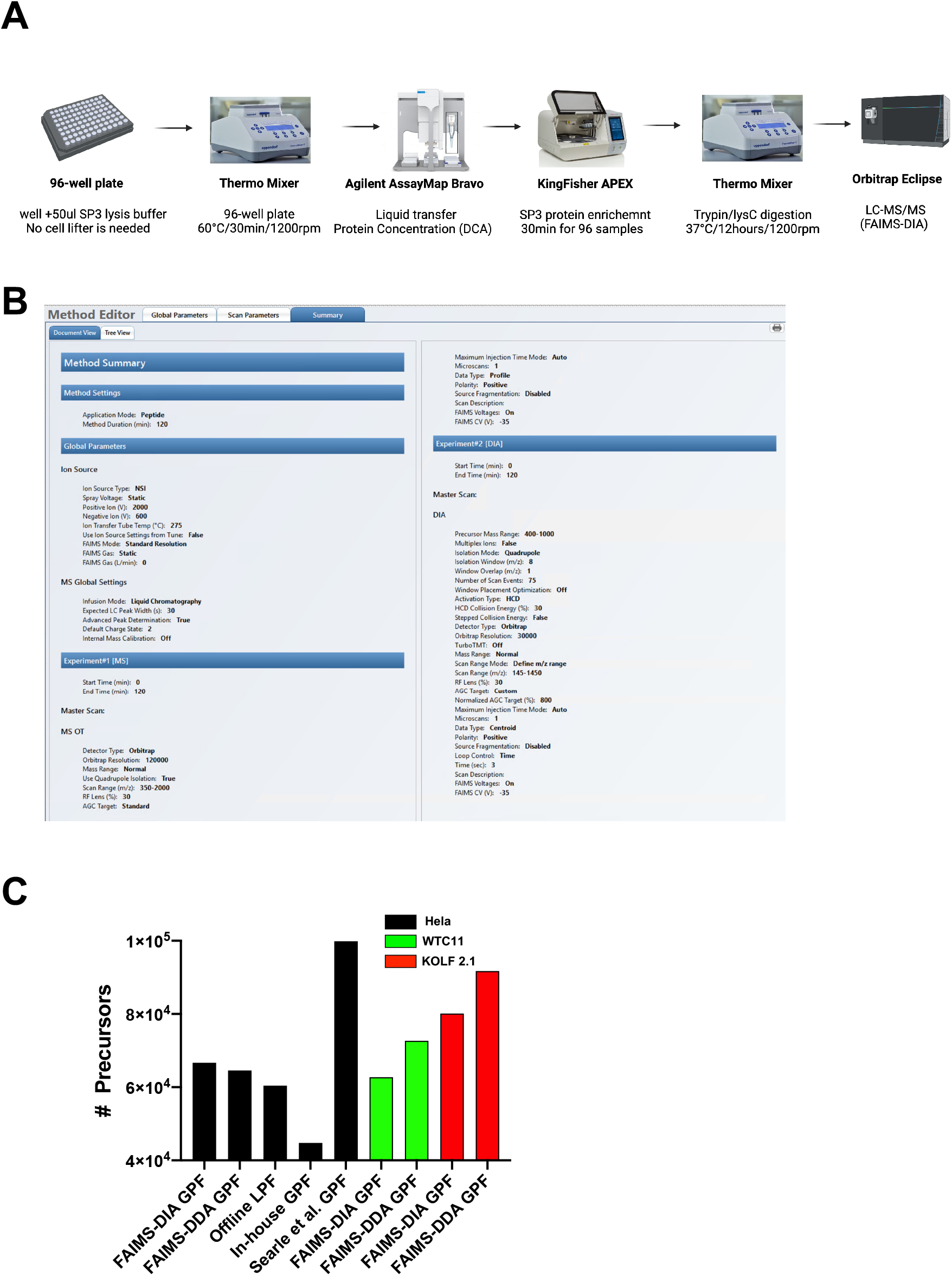
**(A)** Schematic shows a fully automated SP3 sample preparation pipeline. **(B)** Summary of detail MS instrument parameters for FAIMS (−35V)-DIA method. **(C)** Bar chart shows the number of precursor ions in generated DIA libraries.

**Supplemental Fig. S2.**
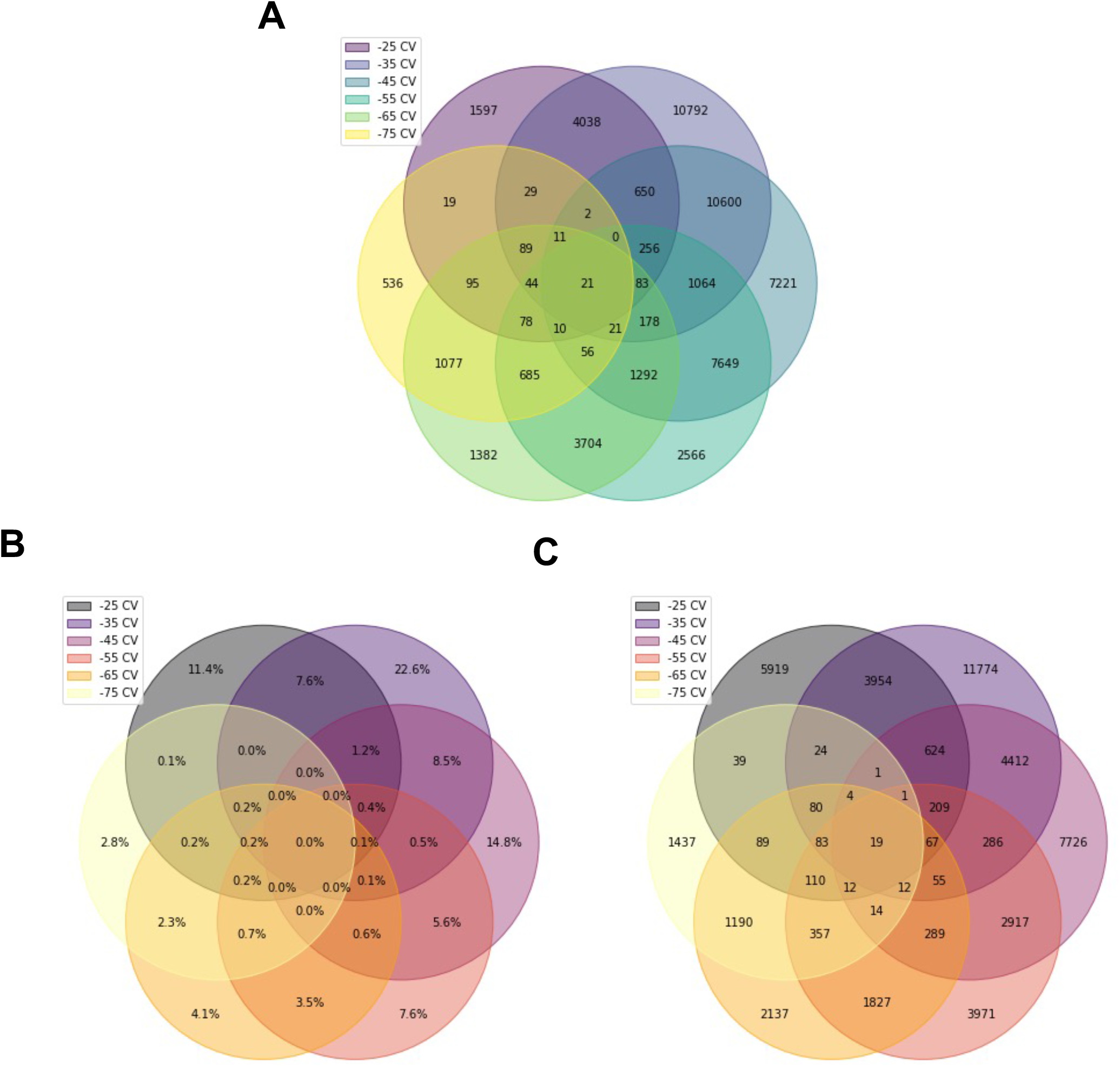
Extension of Fig. 2. **(A)** Venn diagram shows the number of peptides exclusively identified in 6 FAIMS single-CV-based DIA runs (−25V to −75V). **(B-C)** Venn diagram shows the percentage **(B)** and number **(C)** of peptides exclusively identified in 6 FAIMS single-CV-based DDA runs out of all peptides identified in the entire pool (−25V to −75V).

**Supplemental Fig. S3.**
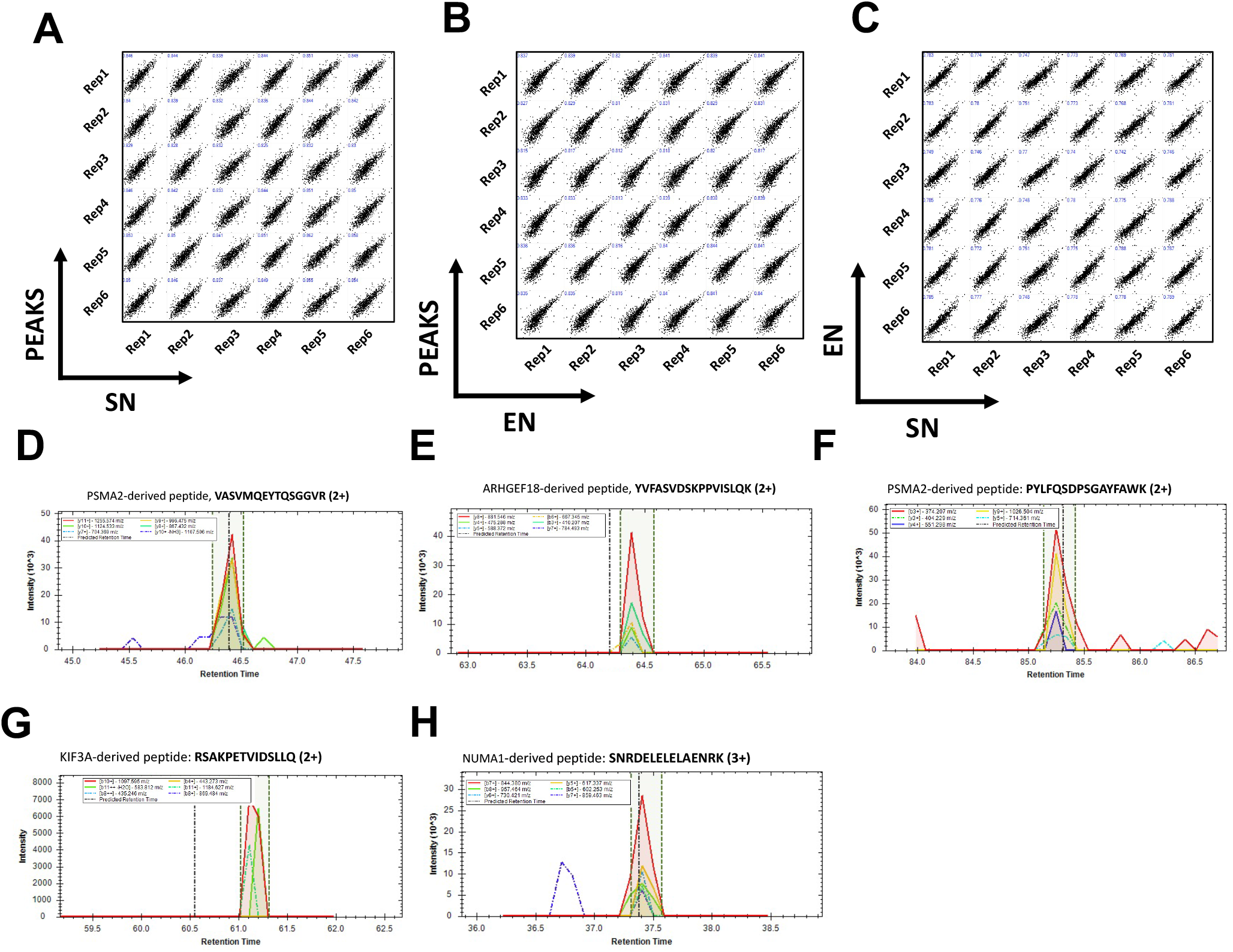
Extension of Fig. 3. **(A-C)** Multi-correlation among protein abundance of 6 biological replicates of KOLF2.1J iPSCs database searched by SN, EN, PEAKS. The correlation between PEAKS and SN is shown in **(A**), PEAKS and EN is shown in **(B)**, EN and SN is shown in **(C)**. **(D-H)** Representative MS2 transitions of peptides only identified by SN, which are YVFASVDSKPPVISLQK **(D)**, VASVMQEYTQSGGVR **(E)**, PYLFQSDPSGAYFAWK **(F)**, RSAKPETVIDSLLQ **(G)**, and SNRDELELELAENRK **(H)**.

**Supplemental Fig. S4.**
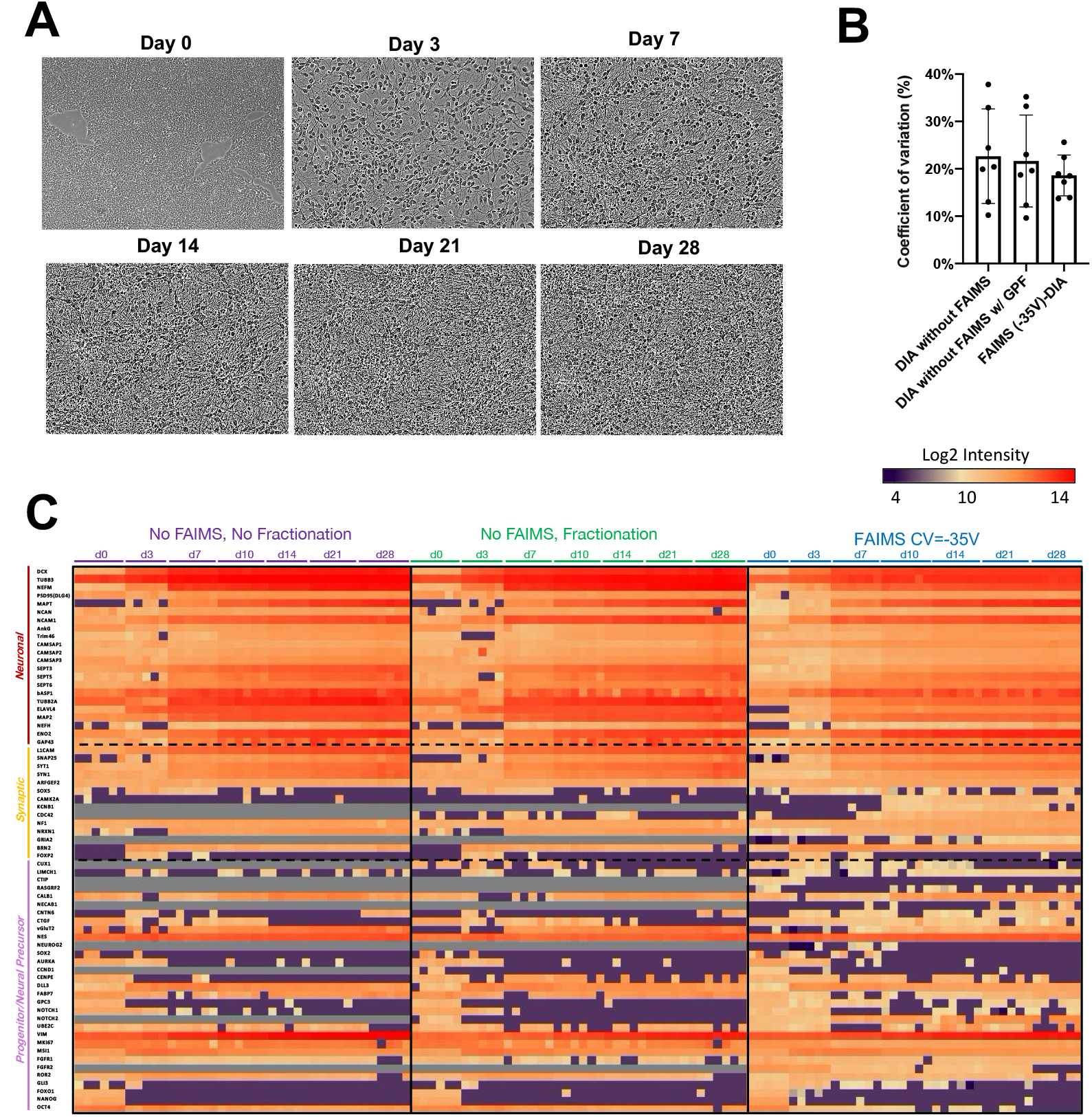
Extension of Fig. 4. **(A)** Representative live cell images of cell morphology of iPSC (d0), i3 Neurons d3, d7, d14, d21, d28. **(B)** Coefficient of variation % (CV%) of identified proteins by different direct-DIA approaches: noFAIMS, noFAIMS with GPF, FAIMS CV=-35V. **(C)** heatmap shows the expression partners of known neuronal, synaptic, and neural precursor markers during neuron differentiation; data points of individual replicate are shown.

**Supplemental Fig. S5.**
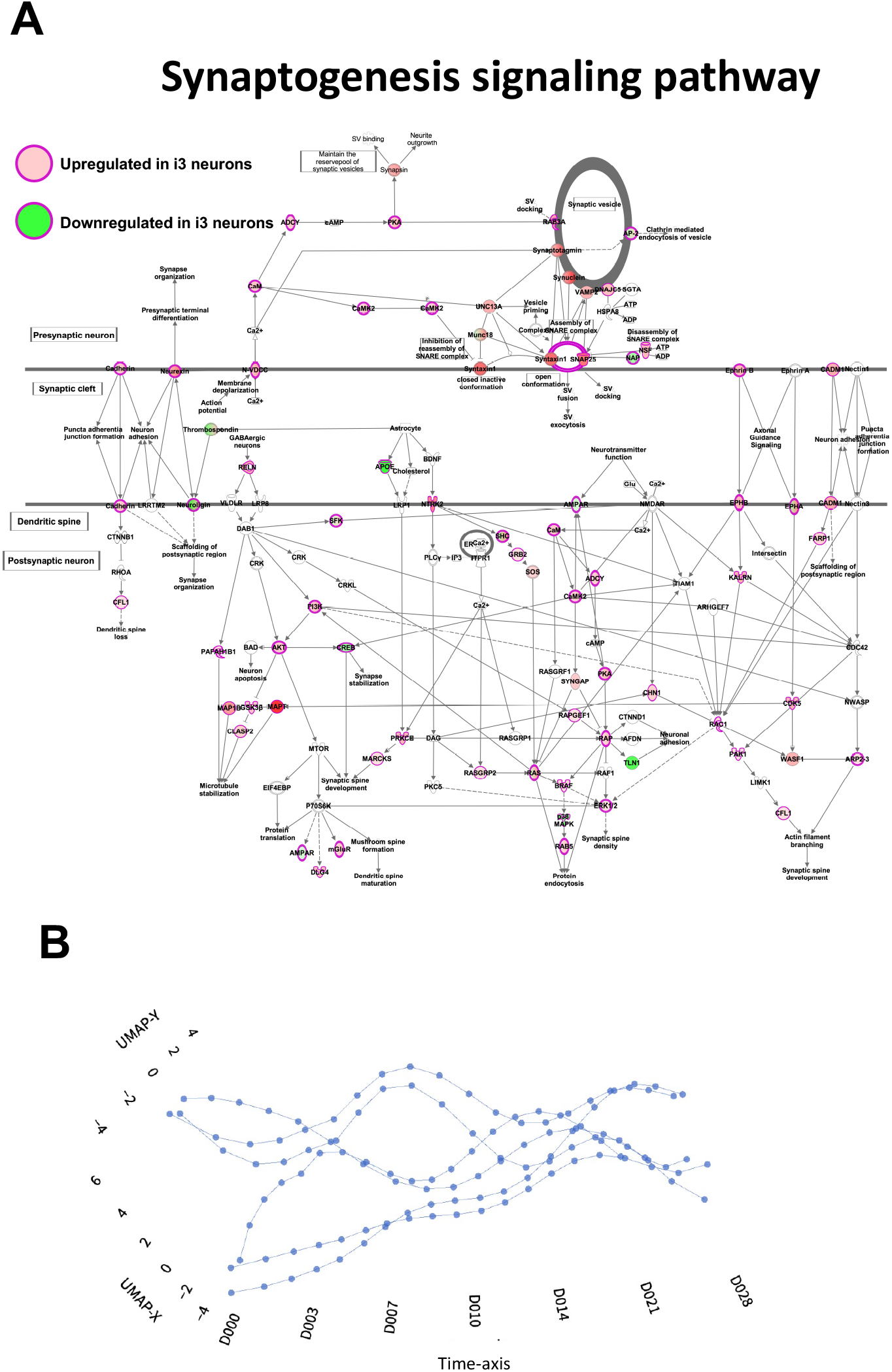
Extension of Fig. 5. **(A)** Synaptogenesis signaling pathway from integrated pathway analysis is shown, red-colored nodes are upregulated proteins whereas green-colored nodes are downregulated proteins identified in current study. **(B)** The shown visualization plots are generated for optimal hyperparameters of Aligned-UMAP using manually inspection. Next section allows to explore the effect of hyperparameters on visualization space. The optimal hyperparameters are metric=euclidean; alignment_regularisation=0.003; alignment_window_size=3; n_neighbors=3; min_dist=0.01; num_cores=32.

**Supplemental Fig. S6.**
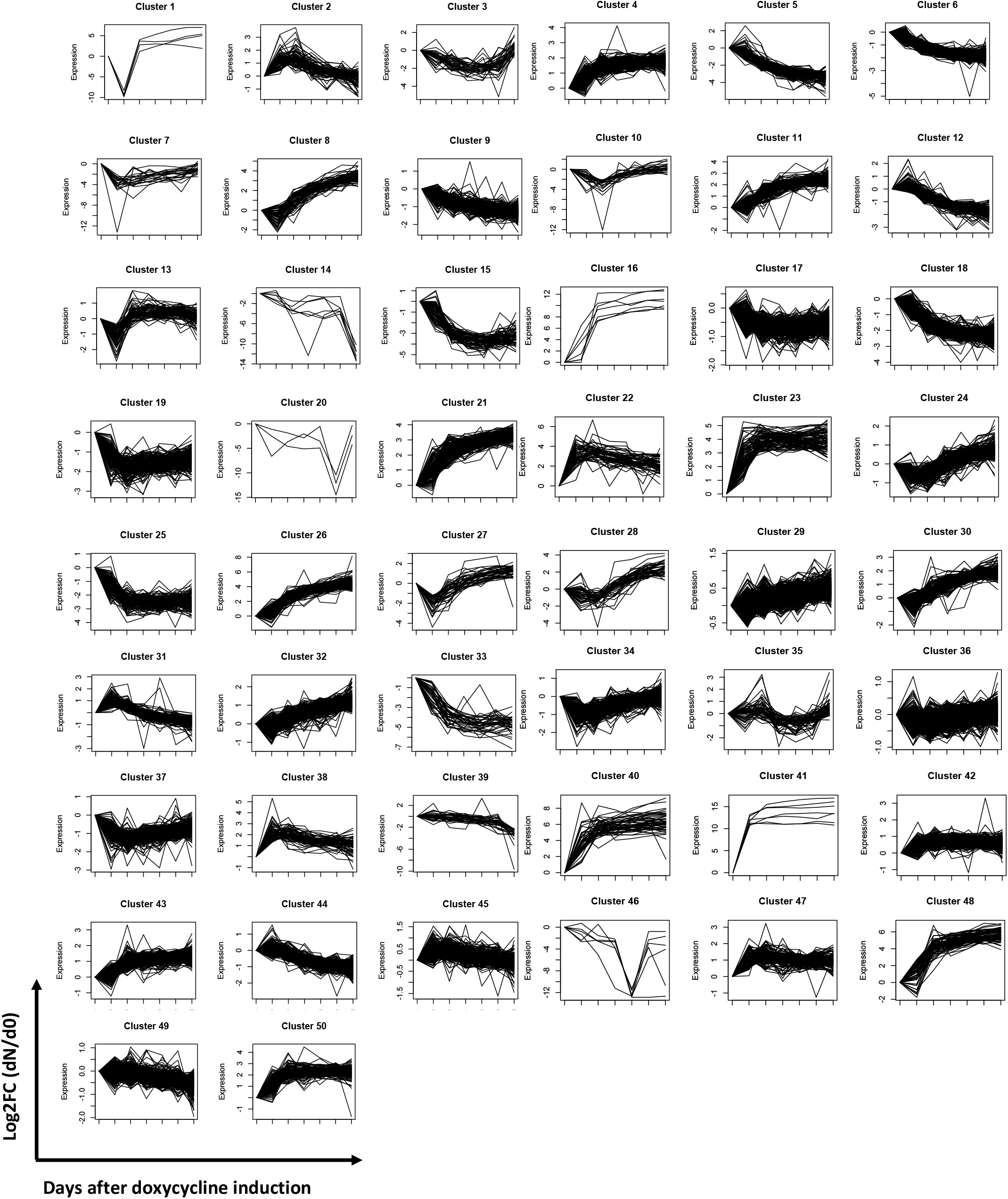
Extension of Fig. 6. All 50 clusters show the protein expression trajectories in the longitudinal proteomic analysis.

**Supplemental Fig. S7.**
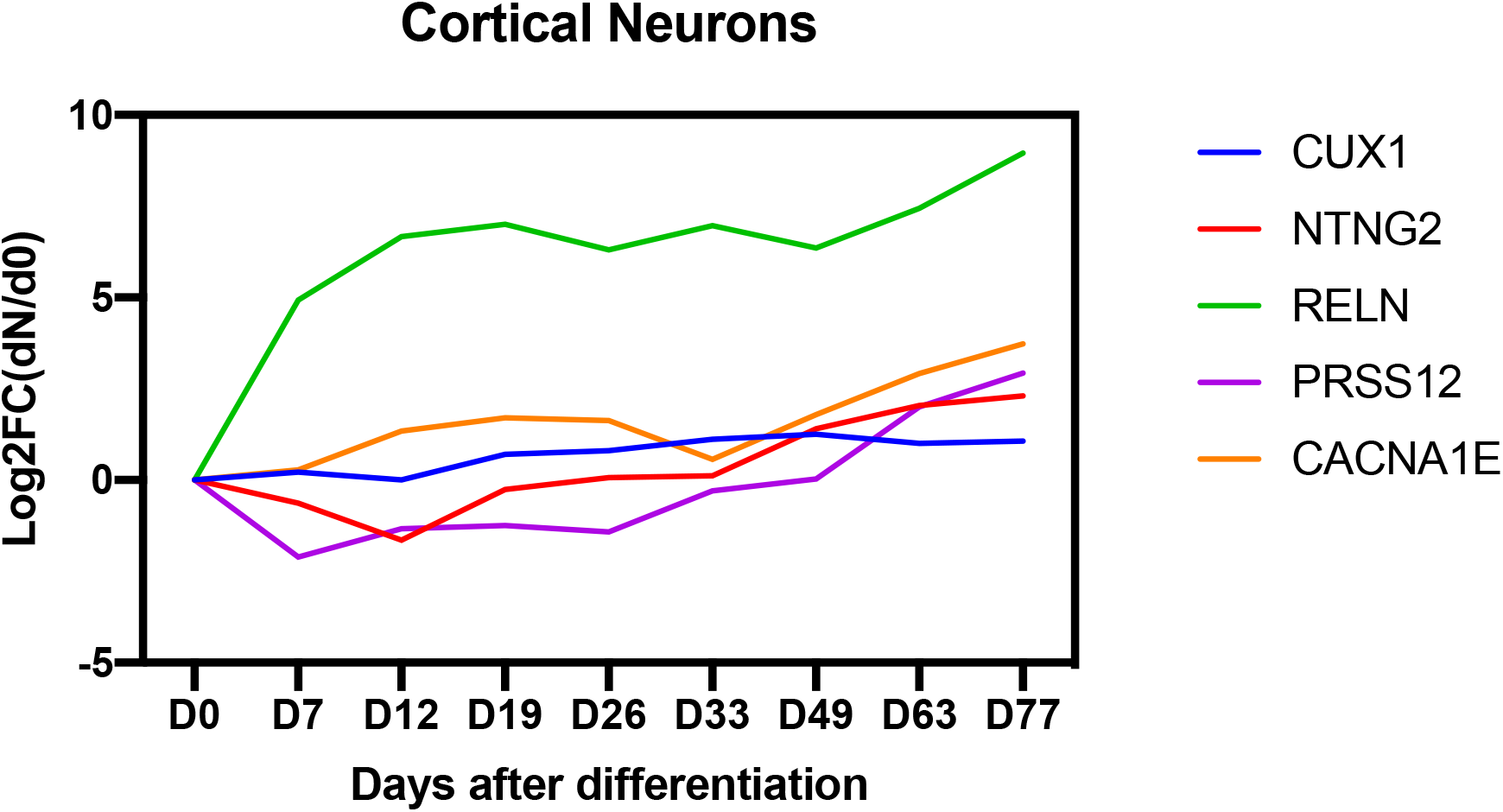
Cortical neuron markers identified in CORTECON, showing their gene expression dynamics over a time course of 77-day differentiation (42).

